# Synthesis of ionizable lipopolymers using split-Ugi reaction for pulmonary delivery of various size RNAs and gene editing

**DOI:** 10.1101/2024.06.11.598497

**Authors:** K. Yu. Vlasova, A. Kerr, N.D. Pennock, A. Jozic, D.K. Sahel, M. Gautam, N.T.V. Murthy, A. Roberts, M.W. Ali, K.D. MacDonald, J. Walker, R. Luxenhofer, G. Sahay

**Author notes:** Both authors contributed equally (K. Yu. Vlasova, A. Kerr).

## Abstract

We present an efficient approach for synthesizing cationic poly(ethylene imine) derivatives using the multicomponent split-Ugi reaction to rapidly create a library of complex functional ionizable lipopolymers. We synthesized a diverse library of 155 polymers, formulated them into polyplexes to establish structure-activity relationships crucial for endosomal escape and efficient transfection. After discovering a lead structure, lipopolymer-lipid hybrid nanoparticles are introduced to preferentially deliver to and elicit effective mRNA transfection in lung endothelium and immune cells, including T cells with low *in vivo* toxicity. The lipopolymer-lipid hybrid nanoparticles showed 300-fold improvement in systemic mRNA delivery to the lung compared to *in vivo*-JetPEI^®^. Lipopolymer-lipid hybrid nanoparticles demonstrated efficient delivery of mRNA-based therapeutics for treatment of two different disease models. Lewis Lung cancer progression was significantly delayed after treatment with loaded IL-12 mRNA in U155@lipids after repeated i.v. administration. Systemic delivery of human CFTR (hCFTR) mRNA resulted in production of functional form of CFTR protein in the lungs. The functionality of hCFTR protein was confirmed by restoration of CFTR-mediated chloride secretion in conductive airway epithelia in CFTR knockout mice after nasal instillation of hCFTR mRNA loaded U155@lipids. We further showed that, U155@lipids nanoparticles can deliver complex CRISPR-Cas9 based RNA cargo to the lung, achieving 5.6 ± 2.4 % gene editing in lung tissue. Moreover, we demonstrated successful PD-1 gene knockout of T cells *in vivo*. Our results highlight a versatile delivery platform for systemic delivering of mRNA of various sizes for gene therapy for a variety of therapeutics.

## Introduction

Rapid advances in mRNA gene therapy and vaccine development require investigation of safe and effective non-viral delivery systems which both protect the mRNA molecules and facilitate their entry into target cells.^1,2^ These carriers are essential to protect the mRNA from degradation in the bloodstream, aid in its cellular uptake, and facilitate mRNA escape from the endosome to allow the mRNA to be translated into functional proteins within the cells. However, the delivery of mRNA of various sizes faces several challenges. For example, larger mRNA molecules are generally more prone to degradation by nucleases and other enzymes, and efficient packaging into a delivery system faces more challenges, which can impact the efficiency of protein expression, and delivery of larger mRNA molecules may be more likely to trigger stronger immune responses compared to smaller ones.^3,4^ Addressing these challenges requires the development of tailored delivery systems and strategies tailored to the specific characteristics of mRNA molecules of various sizes. This includes optimizing nanoparticle formulations and modifying mRNA structures for enhanced stability. Therefore, the design of a universal system that can efficiently encapsulate and deliver mRNA of various sizes or mixtures of mRNAs without requiring additional changes can have a significant impact.

Poly(ethylene imine) (PEI) has been intensively investigated as a cationic polymer for gene delivery applications for decades.^5,6^ However, there is a tradeoff between effective transfection and cytotoxicity for higher molar mass PEIs, which limits its use *in vivo*.^7^ The high charge density of PEI is believed to lead to strong interactions with cellular membranes, which contributes to its cytotoxicity.^8^ However, the amine units of PEI are amenable to chemical modification. Indeed, chemical modifications, or partial hydrolysis and partial reduction^9^ of a poly(2-oxazoline) precursor,^10^ can serve to lower the charge density thus reducing toxic effects^6,11,12^, and modulating polyplex properties, increasing transfection efficacy. For example, inclusion of hydrophobic groups onto polycations has been shown to strongly influence transfection,^13^ presumably due to strengthening self-assembly behavior of the polyplex and modulation of polyplex-cell interactions.^14,15^. Specifically, alkylation of primary amines in branched PEI with dodecyl chains was shown to improve transfection 5- fold while lowering toxicity.^16^ Modification of a low molecular weight PEI (1.8 kg mol^-1^) with various methylcarboxytrimethylene carbonate derivatives showed improved transfection for ethyl and benzyl side chain substituents.^17^ It was found that moderate degrees of modification of branched PEI with propionic acid improved transfection, demonstrating the importance of a balance of hydrophobic moieties.^18^ A range of other promising modifications have been identified, including succination,^19^ acetylation,^20^ carbamoylation,^21^ as well as conjugation with dexamethasone,^22^ lipids,^23–26^ fluoroalkanes,^27^ and aromatic groups,^28^ *inter alia*.^29–33^

While diverse libraries of cationic polymers are being synthesized to enable delivery across extrahepatic tissues, the therapeutic index remains limited for clinical development.^34–37^ Moreover, the dispersity inherent to polymers presents further challenges for scale up and reproducibility. To this end, we hypothesized that a rapid high-throughput method of derivatization of PEI that results in unprecedent structural diversity of PEI based ionizable lipopolymers and further formulation optimization can result in resolving these long-standing issues.

The Ugi multicomponent reaction^38^ is one of best known and versatile multicomponent reactions and utilizes four different reagents - an amine, aldehyde, isocyanide and carboxylic acid - ^39^ and has been utilized in polymer science as a step growth polymerization reaction,^40–42^ a post-polymerization modification tool^43^ and as a polymer coupling reaction.^44^ Here, we introduce the so-called split-Ugi,^45,46^ to synthesize linear lipo-PEI derivatives optimized for RNA transfection. The secondary amine units of linear PEI require this variant of the Ugi reaction, effectively involving two equivalents of secondary amine as opposed to one equivalent of a primary amine in the standard Ugi reaction. The split-Ugi modification is expected to yield a product with two modified repeat units, one a tertiary amine with one substituent stemming from the isocyanide and the one from the aldehyde, and the other an amide group featuring the carboxylic acid moiety (Fig. 1). This introduces a convenient route to prepare novel PEI derivative libraries with a mixture of large variety of functional groups to explore the effect of polymeric structure on transfection efficiency both *in vitro* and *in vivo*. In this study, a library of isocyanide/aldehyde/carboxylic acid reagents were selected to create a range of structures, and additionally molecular weights and modification densities of the PEIs samples were varied to further increase structural diversity (Fig. 1). Through initial screening we identified a lead lipo-PEI with enhanced endosomal escape, subsequent *in vitro* transfection and developed hybrid lipopolymer-lipid nanoparticle formulations for *in-vivo* mRNA-based gene delivery. We demonstrate a multiple order increase of *in vivo* mRNA delivery to the lungs via systemic administration compared to the gold standard *in vivo*- JetPEI^®^. The produced system can deliver both small (IL-12 mRNA) and large (hCFTR) functional mRNA preserving expression of therapeutic proteins. Importantly, the delivery system showed efficient gene editing in lungs endothelial and T cells by delivery of Cre mRNA and CRISPR-Cas9 mRNA system with minimal toxicity.

**Fig. 1.**
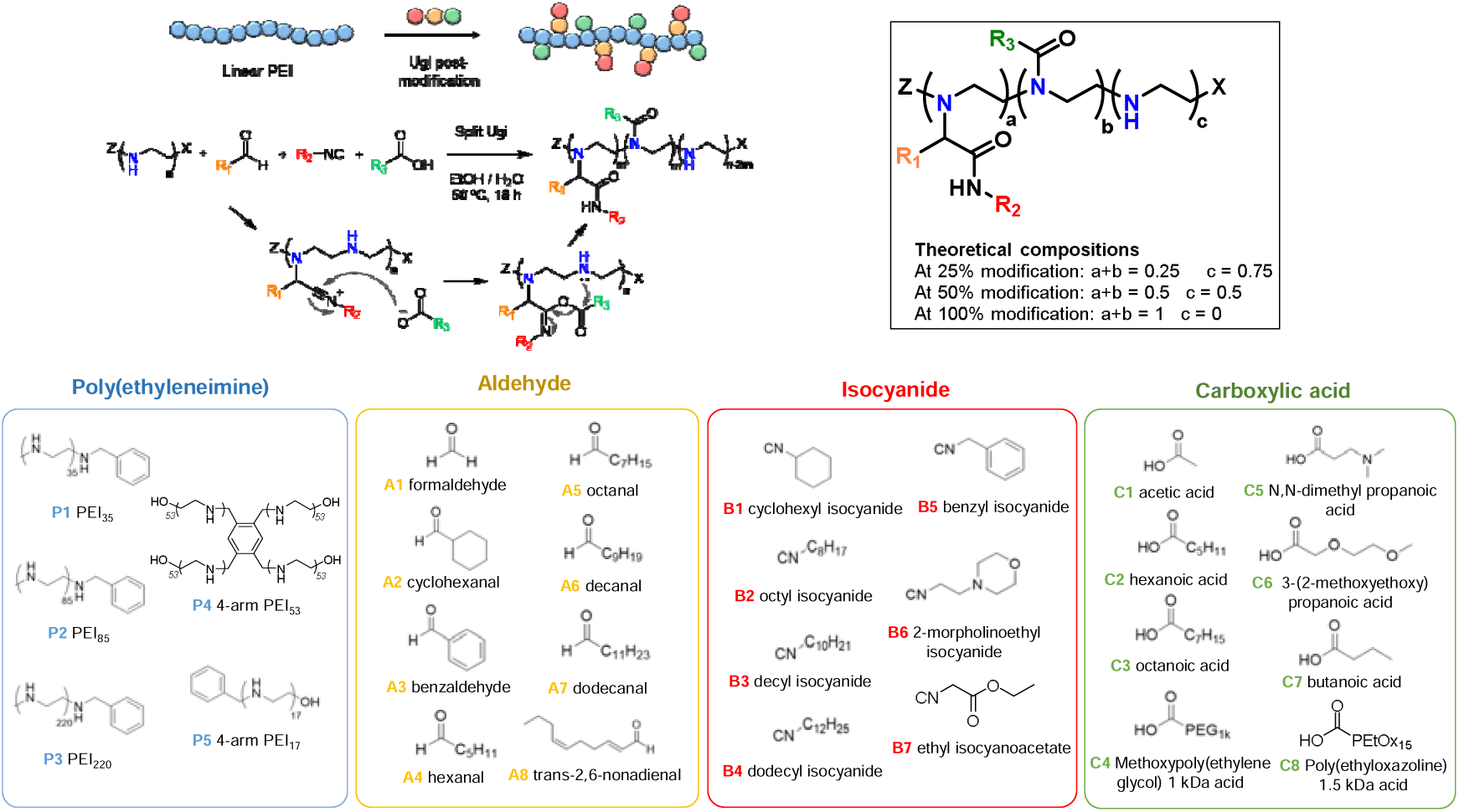
Polymers synthesis and library: scheme for modification of PEI and proposed mechanism of the split-Ugi type reaction; reagent library used in the preparation of PEI derivatives. The structures of molecules are in supporting information Table S1.

## Results

### Synthesis of modified PEI for mRNA delivery

The split-Ugi reaction^47^ is a sparingly employed version of the four component Ugi reaction^48^ that, has not been utilized for polymer modification. We initially assessed the split-Ugi reaction to obtain linear PEI derivatives with a range of reagents and of different modification densities to introduce lipid-like side chains featuring ionizable tertiary amines (Fig. 1). A linear PEI_35_ was reacted with formaldehyde (A_1_), acetic acid (C_1_) and four different isocyanides (B_1_, B_5_, B_6_ and B_7_) targeting a total modification of 50 % of the secondary amine repeat units (4:1:1:1 molar ratio of amine:aldehyde:isocyanide:acid). ^1^H-NMR and ^13^C-NMR spectroscopic analysis showed successful integration of the isocyanide functionality (supporting information Fig. S1), however, the degree of functionalization varied and was lower than expected. In contrast to classic Ugi reactions (utilizing a 1:1 molar ratio of amine to carboxylic acid, or 2:1 for a split-Ugi), we only targeted partial conversion of the amine, requiring an excess of amine with respect to carboxylic acid. We hypothesize that the excess of amine may lower the reactivity of the carboxylic acid in the Ugi by formation of ion pairs. Accordingly, use of excess of the carboxylic acid up to an equimolar quantity with respect to the amine resulted in higher incorporation of all reagents into the polymer for a test system using A_1_/B_1_/C_1_ reagents (Fig. 1). With this, degrees of functionalization close the target was achieved for all the aldehyde and isocyanide components, and therefore this procedure was selected for further derivative synthesis.

An initial library of 148 split-Ugi modified polymers was synthesized by this combinatorial approach, utilizing linear PEI of three different molecular weights (1.6, 3.8 and 9.6 kg mol^-1^), and a 4 arm-star PEI, targeting two modification degrees (25 % and 50 %) with a small library of aldehydes, isocyanides and carboxylic acids (Fig. 1, supporting information Table S1). Linear PEIs were chosen as the polymer backbone, as these can be readily synthesized with low dispersity and well-controlled molar masses using cationic ring opening polymerization of 2-oxazoline and subsequent hydrolysis. Additionally, they are easier to characterize without the complication of mixed primary/secondary/tertiary amine species found in hyperbranched PEIs, which are otherwise preferred in polyplex formation and display the highest transfection.^49^

Successful incorporation of the aldehyde/isocyanide functional groups to a degree matching the targeted modification density was confirmed with NMR spectroscopy. However, the carboxylic acid reagent showed variable incorporation across samples, in some cases we found as low as ∼40 % of the expected amide group formation (supporting information Fig. S1). This leads to a larger fraction of unreacted PEI units, although formation of other side products, unidentifiable in the NMR, cannot be completely ruled out. The incomplete incorporation of carboxylic acid reagent suggests that the imino-anhydride intermediate is not exclusively attacked by the secondary amine as expected in the modified Mumm rearrangement step but may react with another nucleophile in the system. This has been reported when using methanol as the Ugi reaction solvent.^47^ Also the interception by water is known to occur in the 3 component-Ugi reaction, but this typically requires another catalyst and is considered unlikely to occur in the present system. While a range of carboxylic acid reagents was explored, it was not considered to be of high relevance for structural variation due to the lowered incorporation of this reagent onto the polymer. Additionally, preliminary screening of acetic acid derivatives showed promising transfection, which is why acetic acid was used for most of the library. Size-exclusion chromatography (SEC) analysis indicated molecular weights in the expected range and monomodal distributions suggesting the modification otherwise proceeded smoothly (supporting information Fig. S1b).

### Degree of polymerization and length of carbon tail chain effects mRNA delivery

Polyplexes are widely investigated non-viral type of gene delivery system, which provide protection of mRNA and facilitates the endosomal escape. Therefore, for an initial screening we formulated polyplexes from the synthesized PEI-derivatives library via an ethanol injection method.

For *in vitro* studies a library of polyplexes with mRNA encoding Firefly luciferase (Fluc) was screened in HeLa cells to screen mRNA transfection. Polyplexes with mCherry RNA were tested in Gal9-HEK293 cells to evaluate the endosomal escape efficacy as it was described previously.^50^ The endosome damage results in redistribution of Gal9 and is visualized as GFP puncta in the Gal9-GFP reporter cells. Based on *in vitro* screening (Fig. 2a), several trends were observed: first, the more hydrophobic samples appeared to perform better (e.g., U12, U15, U22, U46 (more hydrophobic) compared to U1-U8 (less hydrophobic)). Hydrophobicity of the polymers was varied with a combination of alkyl chain bearing isocyanide and aldehyde reagents. Such functionalities result in a polymer repeat unit with an ionizable tertiary amine unit in the backbone and two alkyl chain groups attached, reminiscent of cationic lipids, which are known as excellent transfection agents (e.g., DOTAP, DOTMA or DOGS).^51^ Notably, the position of hydrophobic chain affects the transfection rates as well.

**Fig. 2.**
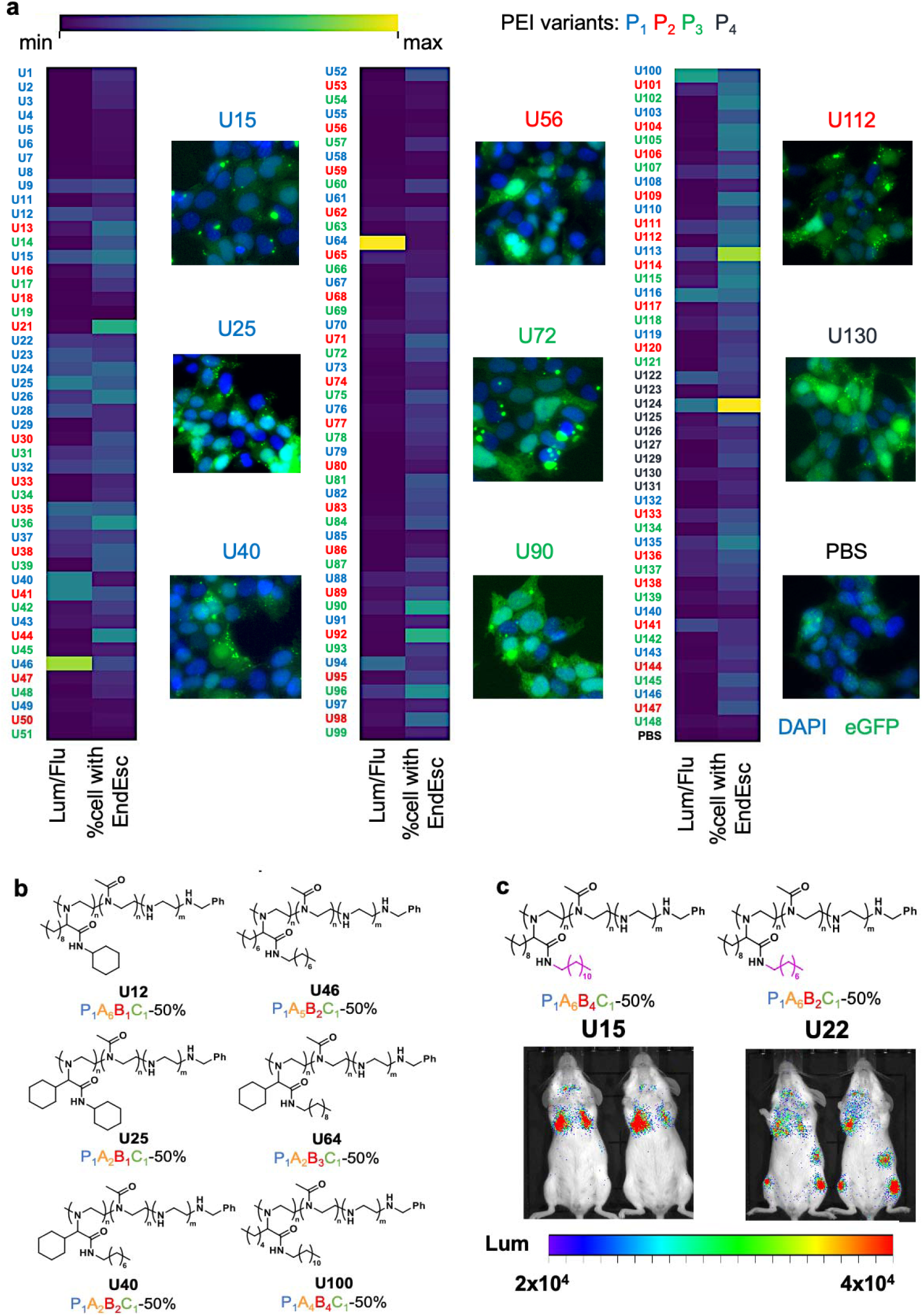
Polymers *in vitro* and *in vivo* screening. **a)** *In vitro* library screening. HeLa cells were treated with Fluc mRNA-loaded lipopolyplexes. The relative luciferase expression (normalized to cell viability fluorescent signal) after 24 h incubation with lipopolyplexes is shown on the heat map (n=6-8). Gal9-HEK293 cells were treated with mCherry RNA-loaded lipopolyplexes. The percent of cells with endosomal escape (EndEsc) cases after 24 h incubation with polyplexes is presented on the heat map (n=40-150). Data are presented as Means. Representative images of Gal9-HEK293 cells after treatment are shown; **b)** structures of *in vitro* high performing lipopolymers; **c)** *in vivo* screening of grouped Fluc mRNA-loaded lipopolyplexes. Representative IVIS images of BALB/c mice 5-6 h following polyplexes administration. For *in vivo* screening studies, results were obtained from two mice per group. Each particles type in the group was injected at the dose of 10 μg Fluc mRNA/mouse.

Thus, variation of carboxylic acid substitutions showed that the shortest alkyl chain (acetic acid) is the most efficient of those investigated. Additionally, the lower molar mass polymers and the higher modification density showed better performance. For PEI homopolymers, higher molar mass show higher transfection, however, this trend does not apply for the hydrophobic derivatives in our library. The addition of hydrophobic interactions is expected to strengthen interactions of the polyplex assembly, leading to more stable particles for even the low molar mass derivatives.

Next, a selection of these polyplexes was screened *in vivo*. To accelerate screening, we pooled formulations, encapsulating Fluc mRNA, into groups with 15-22 individual polyplexes per group (1 μg dose of Fluc mRNA each) and injected them intravenously (i.v.), which reduced the number of animals and the time required to identify lead polyplexes. Neither of the tested groups showed *in vivo* transfection activity (supporting information Fig. S2), nor did they show activity upon intramuscular (i.m.) injection (supporting information Fig. S2). Only after intraperitoneal (i.p.) injection, weak transfection signals were detected in lungs. Nevertheless, we considered that the produced polyplexes might work *in vivo* at higher doses, as does the commercial *in vivo* JetPEI^®^, and stability to aggregation or dissociation is crucial parameter. It is worth noting, low animal activity and ruffled fur were observed after injections of some groups, indicating potential inflammation. Polymers such as U12, U 25, U40, U46, U64 and U100 (Fig. 2b), which performed well *in vitro,* produced large particles unstable upon storage as judged by dynamic light scattering analysis (supporting information Size screening). Thus, moving to higher doses, we only injected (i.v.) polyplexes made using polymers U15 and U22, which produced stable formulations at high concentrations and showed *in vivo* activity at dose 10 μg Fluc mRNA per mouse. Both formulations showed weak transfection signal. Notably, U15 with longer acyl chain of isocyanide substitute (CN-C_12_H_25_) compared to U22 (CN- C_8_H_17_) showed evident accumulation and Fluc mRNA transfection in lungs while U22 polyplex distributed Fluc mRNA mostly to the lymph nodes and spleen (Fig. 2c). We assume this could be due to higher hydrophobicity of U15 and, as a result, better stability of particles, which prevents polyplexes from aggregation or dissociation in the presence of negatively charged blood components.^52^ Moving forward, we decided to further iterate on the polymer structure basing from the hit compound U15.

Our *in vitro* screening of the initial library revealed that most of polymers facilitate the endosomal escape, but not mRNA transfection (Fig. 2a). The reason for low transfection rates with such type of materials could be a limited cargo release after endosomal escape, which in turn could be correlated to the molar mass of the polymer derivatives. Therefore, the polymeric library was expanded with U15 analogs of lower molar mass of the starting PEI (880 g mol^-1^), while targeting higher modifications of 66% (U154) and 100 % (U155), to improve the performance of polyplexes *in vivo*. *In vitro* screening (Fig. 3a, b) of these polymers showed the following patterns: the mRNA cargo release is reduced with increasing molar mass of the polymer (U15-U16-U17, respectively), presumably due to their elevated stability and profound interactions with mRNA. At the same time, as the molar mass of polymer was reduced significantly (U154 compared to U15), transfection was also reduced due to lower particles stability. The preference of higher modification density (U155 compared to U154) may suggest a reduction in charge density is beneficial, due to the larger number of non-ionizable amide repeat units introduced. Additionally, the presence of sufficiently hydrophobic groups appears to be necessary for particle stability and optimal transfection. It should be noted that the split-Ugi not only transforms secondary amine to an amide, it transforms secondary amine into a tertiary amine.

**Fig. 3.**
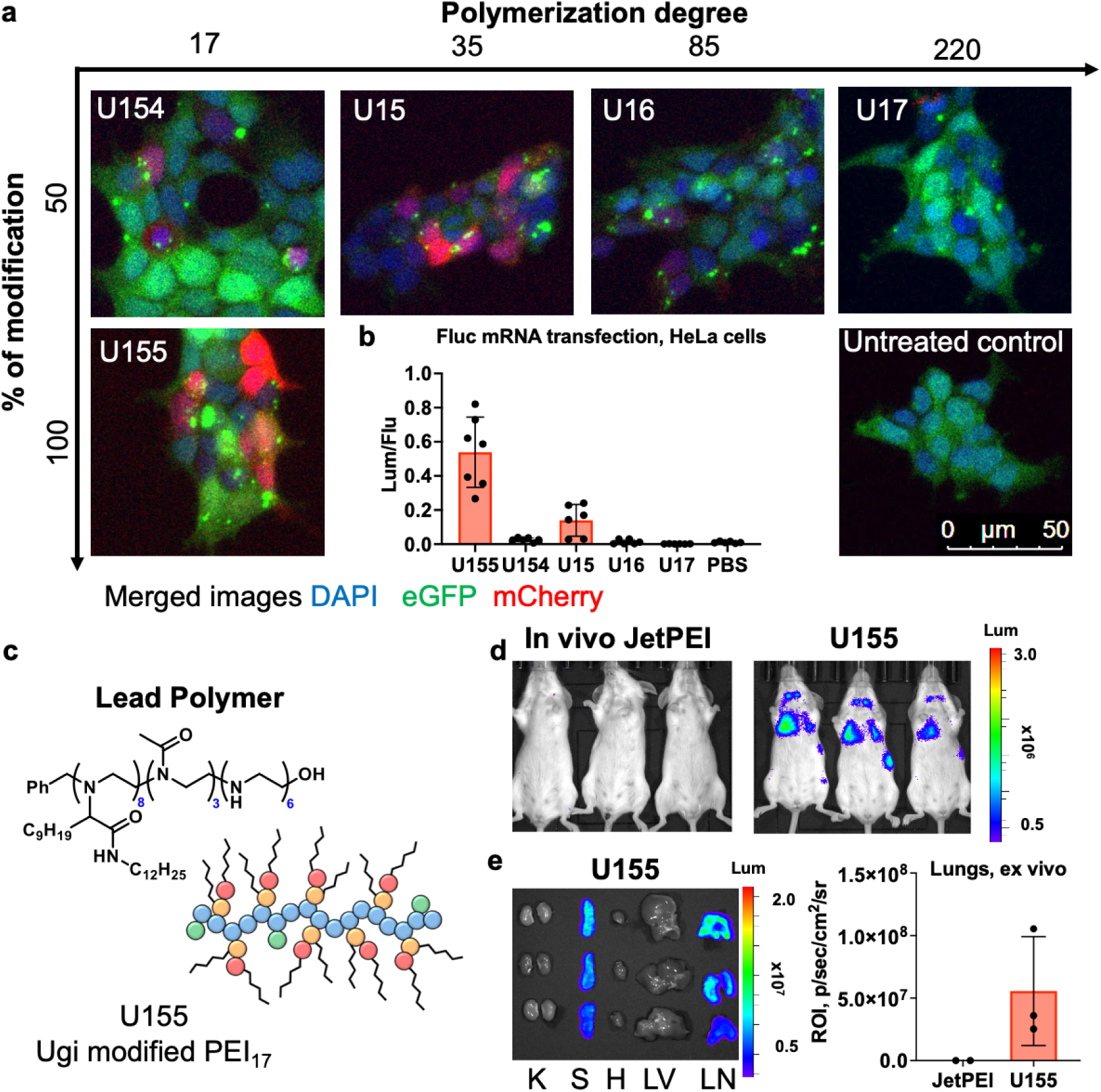
*In vitro* and *in vivo* evaluation of lipopolymers. **a)** Representative images of Gal9-HEK293 cells after 24 h incubation with mCherry RNA-loaded lipopolyplexes from polymers of various chain length and modification; **b)** HeLa cells were treated with Fluc mRNA-loaded lipopolyplexes. The relative luciferase expression (normalized to cell viability fluorescent signal) after 24 h incubation with polyplexes is shown. Data are presented as Mean±SD (n=6-8); **c)** Structure of lead lipopolymer U155; **d)** *In vivo* and **e)** *ex vivo* bioluminescent images of BALB/c mice 5-6 h post i.v. injection of 5 μg Fluc mRNA per mouse. K- kidney, S – spleen, H – heart, LV – liver, LN – lungs. Data are presented as Mean±SD (n=2-5).

Thus, a new lead polymer with promising transfection performance was identified (U155), synthesized by modification of a PEI_17_ with decanal, dodecylisocyanide and acetic acid reagents via the split-Ugi (P_5_, A_5_, B_4_, C_1_) (Fig. 3c). The targeted 100 % modification should yield a polymer with 50% tertiary amine repeat units featuring a lipid-like tail, and by ^1^H NMR a functionalization of 47 % = (supporting information Fig. S1c,d). Notably, the number of methyl amide repeat units formed was substantially lower at 19 % (target 50 %), meaning a significant portion of secondary amine PEI units remain despite targeting full conversion.

We next explored the capability of the selected polymer for *in vivo* Fluc mRNA delivery. Formulated by vortexing, polyplexes exhibited a diameter of 201±41 nm (PDI 0.20±0.05) and the ζ potential of 57±6 mV, as was characterized by dynamic light scattering (DLS). Injected (i.v.) Fluc mRNA encapsulated into U155 polyplexes showed strong bioluminescent signal and the preferential Fluc protein expression in lungs and spleen (Fig. 3d, e). Notably, the gold standard *in vivo* JetPEI^®^ did not show transfection activity at the same dose (5 μg mRNA per mouse). Accumulation of cationic gene delivery systems in the lungs are well documented and various pathways of nanoparticles fate after i.v. administration are described.^53,54^ One of the possible way of lungs accumulation is through opsonization of proteins like vitronectin^54^ or when highly charged cationic particles interact with cellular blood components, especially with erythrocytes, producing small aggregates which firstly infiltrate in fine lung capillaries and then translocate to the spleen and liver.^53^ Additionally, biodistribution to the lungs and spleen has been found to be highly dependent on particles size.^55^ Due to the high hydrophobicity of the polymer, U155 polyplexes are prone to aggregation during the concentration process, which resulted in the formation of particles in the range of >400 nm. This characteristic increases their endocytosis by splenocytes (supporting information Fig. S3d). Overall, the highly positively charged U155 polyplexes are rapidly eliminated from the circulation within 24 h (supporting information Fig. S3a-c). To prevent the described processes, we assumed that a reduction of the particle charge, size stabilization and particles surface protection could be beneficial. To do so, we investigated a polyplex-lipid nanoparticle hybrid strategy.

### Hybrid polymer-lipid U155@lipids nanoparticles outperform *in vivo* JetPEI^®^ in mRNA delivery to the lung

Previously it was shown that combination of polymers with lipids for formulation of gene delivery systems can improve colloidal properties of such particles under physiological conditions, both enhancing target delivery and as well as increase *in vivo* transfection efficiency.^55,56^ To decrease the ζ -potential of resulting hybrid nanoparticles negatively charged phosphatidyl glycerol (PG) lipid can be added.^57^ Pattipeiluhu et al. previously used 1,2-distearoyl-*sn*-glycero-3-phospho-(1’-rac-glycerol) (DSPG) in LNPs to manipulate distribution of particles after i.v. injections.^58^ Inclusion of anionic lipid into LNPs reduces accumulation of particles in liver, while mean positive ζ -potential leads to lungs accumulation.^59,60^ However, anionic LNPs have a rather low RNA loading efficiency, due to the charge repulsion against mRNA molecules.^61^ To overcome this limitation lipopolyplex particles were formed via two-step technique.^62^ firstly, cationic polyplex formation, and secondly addition of anionic lipid membrane. Such composition improves stability of particles and can reduce toxicity. Accordingly, using U155, hybrid polymer-lipid nanoparticles were formulated (Fig. 4a). Briefly, polyplexes formed using the approach described above were mixed (2/1 w/w) with an ethanol solution containing DSPG, soy PC, cholesterol and DMG-PEG2000 (molar ratio 22/23/50/5) using microfluidic mixing (Fig. 4b). The final formulation was dialyzed against Tris-HCl buffer (25 mM, pH 7.4) for 3-4 hours at room temperature. Cholesterol plays a crucial role in enhancing the stability of liposomes and lipid nanoparticles (LNPs), which, in turn, affects their blood clearance.^63,64^ Additionally, cholesterol has been found to enhance the transfection efficiency of RNA.^65^ Studies suggest an optimal molar content of cholesterol typically in the range of 38.5-50%,^66,67^ here, we chose 50% of cholesterol, aligned with literature.^68^ The optimal molar concentration of PEG-lipid in liposomes can vary depending on the specific application. To evade the body’s immune system and extend liposomes circulation time it was shown that 5% molar PEG-lipid is optimal.^69,70^ Moreover, Kaczmarek *et al.* showed that inclusion of 2-5 mol % of C18-PEG2000 in hybrid polymer-lipid nanoparticles is the most efficient for lung-targeting.^55^

**Fig. 4.**
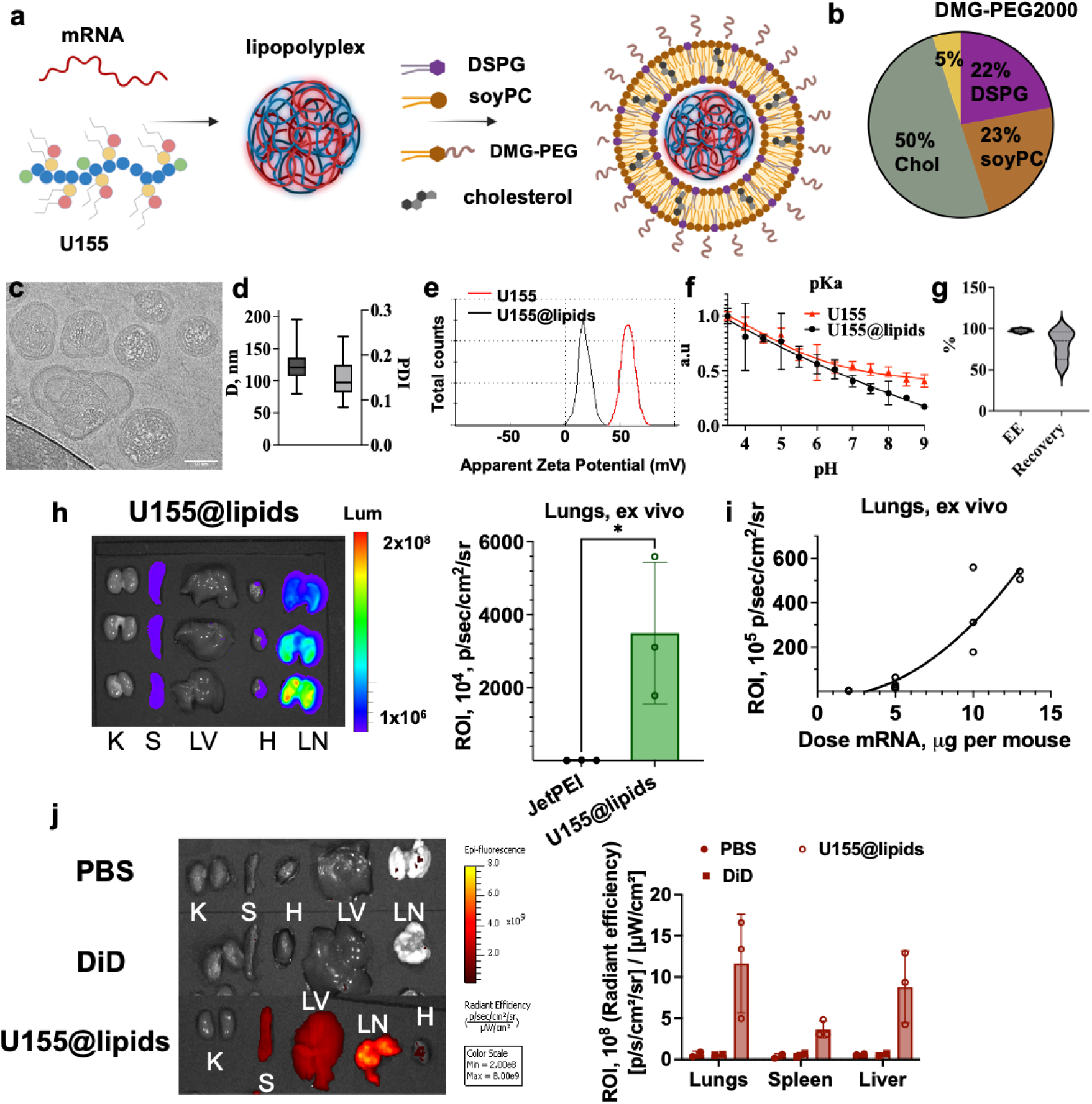
Manufacturing and use of lipopolymer-lipid hybrid nanoparticles. **a)** Schematic illustrating the formulation of a lipopolymer lipid hybrid nanoparticles U155@lipids; **b)** lipid composition of outer shell of U155@lipids; **c)** representative Cryo-TEM image of U155@lipids. Scale bar, 50 nm; **d**) hydrodynamic diameter and PDI of produced nanoparticles in 25 mM Tris-HCl buffer (pH 7.4); **e)** ξ-potential of U155 lipopolyplexes and U155@lipids in 2.5 mM Tris-HCl buffer (pH 7.4); **f)** normalized TNS fluorescence of nanoparticles at various pH. TNS interacts with positively charged amines and produces fluorescence signal; **g)** encapsulation efficiency (EE) and recovery of mRNA after loadin into U155@lipids (n=3); **h)** *ex vivo* images and quantification of bioluminescence in lungs 5-6 h post i.v. injection of 10 μg Fluc mRNA per mouse. K- kidney, S – spleen, H – heart, LV – liver, LN – lungs. Data are presented as Means±SD (n=3), p<0.05; **i)** *in vivo* dose response of Fluc mRNA transfection in lungs (BALB/c mice) 5-6 h post i.v. injection; **j)** representative images and quantification of tissue biodistribution of DiD-labeled U155@lipids nanoparticles. K- kidney, S – spleen, H – heart, LV – liver, LN – lungs. Data are presented as Means ± SD; n = 3. S – spleen, LN – lungs. Results were obtained from 2 (control) and 3 (treated) mice per group and are presented as Means ± SD.

Addition of DSPG showed improved *in vitro* mRNA transfection (supporting information Fig. S4a, S5). Literature suggests optimal levels of DSPG 20-25 mol % of total lipids in liposomal membrane for *in vivo* studies.^71^ Indeed, we found that an increase in DSPG from 22 mol% to 31.5 mol% resulted in a dramatic reduction of mRNA transfection in the lung after i.v. injection (supporting information Fig. S4b). This reduction was due to lower nanoparticles stability, as the negatively charged head groups of DSPG are in proximity and repel each other, destabilizing the bilayer^71^ or due to shrinking the positive charge further. A U155@lipids formulation was used to study efficiency of mRNA delivery to the lung. Cryo-TEM images showed a rather irregular structure of hybrid nanoparticles (Fig. 4c). Mean hydrodynamic diameter was 120 nm with PDI<0.2 (Fig. 4d). While Cryo-TEM showed some clear variability of particle morphology, DLS suggested a rather low dispersity. Covering the U155 polyplex with a lipid bilayer significantly decreased the total particle ζ-potential to 11.4±7.7 mV (Fig. 4e). Titration with fluorescent dye TNS at various pH values showed a slowly falling trend line, which suggest multiple overlapping pK_a_ values in the system (Fig. 4f). Encapsulation efficiency of produced nanoparticles was above 98% with a good recovery (> 75%) (Fig.4g). The lipid shell increased nanoparticles tropism for the lungs 5-fold compared to bare U155 polyplexes (compare Fig. 3e and Fig. 4i). Overall, the optimized particle demonstrated superior effectiveness compared to the commercially available *in vivo* JetPEI^®^ across multiple doses, particularly showing a 300-fold higher efficacy at 10 µg of Fluc mRNA per mouse (Fig. 4h, supporting information Fig. S6). Interestingly, U155@lipids particles showed a non-linear dose response in lungs (Fig.4i). Such effect has been reported in some cases of various non-liver targeted nanoparticles, in particular polymeric ones.^72,73^ For example, in a tumor model, increased target tissue and cell accumulation of nanoparticles was shown through reduction of uptake by phagocytic cells such as macrophages and dendritic cells in the liver and spleen.^73^ Multiple strategies exist to deplete macrophages: pre-injection of clodronate and other agents or blank particles of the same composition as ones with active molecule.^73–76^ As our formulation contains lipids which have tropism to the liver, we first checked the actual biodistribution of U155@lipids by labeling them with fluorescent dye DiD. *In situ*, produced particles were stable in 50% serum at 37 ^°^C for at least 4 h (supporting information Fig. S7). Mice were injected via tail vein with 5 µg Fluc mRNA nanoparticles with 0.1 mol % DiD and U155@lipids preferentially accumulated in the lungs, to a lesser extent in the spleen and liver (Fig. 4j). We posit the low but noticeable liver accumulation may be due to phagocytosis of these lipids by macrophages that are targeted then towards the liver and differentiate into Kupffer cells.^73^ To test this, we injected blank U155@lipids 12 h prior to administration of 5 µg Fluc mRNA nanoparticles (Fig. 5a). The blank particles were prepared using the same method as the mRNA-loaded particles, and their concentration was adjusted to match the 5 µg dose of loaded particles by volume. Indeed, the pretreatment increased the luciferase expression in the lungs ∼2-fold compared to standard scheme (Fig. 5b). In contrast, co-injection of a mixture of blank particles with Fluc mRNA loaded ones did not affect transfection in lungs. Interestingly, a similar effect was observed after multiple dosages of U155@lipids loaded with Fluc mRNA. Since luciferase expression is cleared from the body within 24-48 h after 5 µg Fluc mRNA administration (supporting information Fig. S8), we injected a second dose of the same formulation 48 h after first administration. We observed 2-fold increase in protein expression after the second dose (supporting information Fig. S9). These data suggest that through saturation of macrophages in liver we can further enhance the accumulation of these nanoparticles in the lungs, which also enables multiple dosing of mRNA without compromised transfection efficiency. Additionally, we assessed the possibility of lung inflammation after first dose (pre-treatment), which could be responsible for enhancement of the particles’ accumulation upon secondary dose.^77^ Lung histological samples revealed no statistically significant difference in immune cells infiltration between U155@lipids and PBS injected animals and did not show signs of tissue damage (Fig. 5b-c). No cytokines, involved in acute inflammation, such as IL-1, IL-6 or TNFα were detected in the serum samples taken before the lungs were removed (Fig. 5d-g). In light of the observed results, which demonstrate the delivery and transfection of Fluc mRNA in multiple organs by our platform, the mechanism of nanoparticles distribution could potentially be understood as ‘passive’ organ tropism, a process that enhances the preferential lungs accumulation by the electrostatic attraction of positively charged nanoparticles and negatively charged blood components (proteins, cells).^78^ This effect is likely due to the high local blood flux and the extensive surface area of pulmonary endothelium, facilitating delivery and uptake of mRNA molecules in the lungs.^79^

**Fig. 5.**
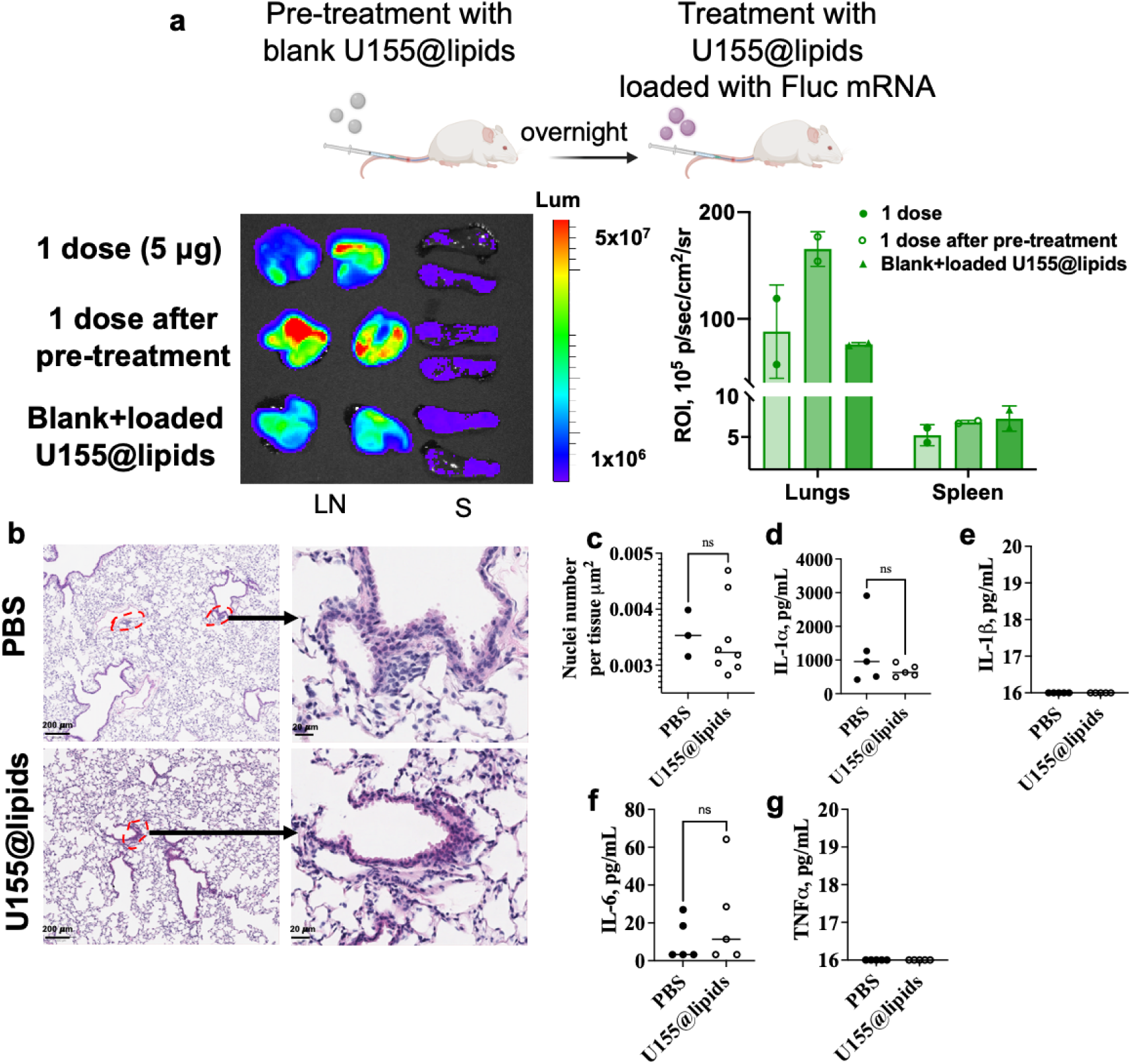
**a)** *Ex vivo* images and quantification of bioluminescence in lungs and spleen 5-6 h post i.v. injection of 5 μg Fluc mRNA (1 dose) per mouse before and after pre-treatment with blank nanoparticles (dose were adjusted by volume compared to loaded U155@lipids). S – spleen, LN – lungs. Results were obtained from two mice per group and are presented as Means ± SD; **b-g)** Inflammation level in lungs 24 h post i.v. injection of 5 μg Fluc mRNA per mouse: **b)** representative images of paraffin-embedded lung sections, which were stained with H&E. Red dashed loops indicate cells infiltration and occurrence of possible micro-abscesses; scale bar 200 μm (left panel) and 20 μm (right panel); **c)** QuPath software quantification of the number of nuclei normalized to tissue area (n = 3 slides for PBS control and n= 8 slides for U155@lipids). Representative image of how QuPath’s tool selects nuclei can be found in supporting information Fig. S10. **d-g)** Plasma concentrations of cytokines IL-1, TNF were below the detection threshold (<16 pg/mL) (n=5 biological replicates).

### Hybrid U155@lipids enable mRNA delivery to the lungs and T cells

To identify the potential applications of U155@lipids, we determined the cell populations within the lungs that express the mRNA product using the Ai9 mouse strain. Upon administration and functional expression of Cre-recombinase, a stop codon is excised from atdTomato protein expression DNA construct present in the ROSA26 safe harbor locus (Fig. 6a). We checked the organ distribution of tdTomato protein by flow cytometry and IHC. First, we collected lungs, spleen and liver and identified the number of tdTomato positive cells by single cell dissociation and using flow cytometry. The lungs were the primary organ that showed significant transfection (Fig. 6b), while the spleen and liver exhibited Cre mRNA transfection to a lower degree (Fig. 6a). These observed results further support the hypothesis of ‘passive’ tissue targeting mechanism by U155@lipids. The majority of tdTomato+ cells in the lungs transfected by U155@lipids were identified as CD31+ (putative endothelial cells) as well as CD45+( immune) cells (Fig. 6b). While CD31+ is commonly used to identify endothelial cells both CD45+ and CD45- cells have been reported to express CD31+. We found that less than 30% of tdTomato+ cells are CD45-/CD31+ endothelial and more than 50% of them are CD45+/CD31+ hematopoietic cells or endothelial precursor cells (Fig. 6b).). Given that our tissue digestion protocol for single cell digests favors immune cell isolation and survival over others cell types,^81^formalin fixed paraffin embedded tissue samples stained with key markers (CD31, CD45 and E-cadherin). Processing of mIHC images of tdTomato+ stained tissue revealed higher transfection rates (Fig. 6d) compared to flow cytometry (Fig. 6b). Multiplex IHC images showed colocalization of tdTomato signal with both endothelial and immune cells (Fig. 6c), however, by IHC CD45-/CD31+ cells were chiefly responsible for the strong tdTomato signal. Notably, based on tissue morphology, U155@lipids nanoparticles not only reach blood vessels, but also Lyve1+ lymphatic vessels (Fig. 6c). Positively charged nanoparticles have been reported to extravasate out of the blood vessel into lymphatics through fenestrations in the endothelium or through transcellular transport through increased interaction and adsorption to the negatively charged cell membranes of endothelial cells.^80,81^ Some rare colocalizations with E-cadherin+ epithelial cells were also observed, while no transfection of large airways was found (Fig. 6c). Among CD45+/tdTomato+ immune cells a portion of T cell receptor positive (TCRb+) (∼5%) and B220+ cells (∼1%) were transfected (Fig. 6b). T lymphocytes are critical regulators and effectors of adaptive immune response, and their transfection *in vivo* offers unique opportunities to advance cancer immunotherapy, autoimmune diseases treatment or vaccine development. Aside from observed transfection in the lung it was noted that a small portion of TCRb+/tdTomato+ cells (∼0.4%) was detected in spleen tissue as well (supporting information Fig. S12). Activation of T cells could be associated with inflammation, however, H&E of lung (Fig. 6c) and liver (supporting information Fig. S14) tissues 3 days post Cre mRNA injection did not reveal any signs of inflammation or tissue damage. Periodic Acid-Schiff (PAS) staining combined with Fast Green counterstaining, which is sensitive to liver function, also did not show evidence of toxicity (supporting information Fig.S14).

**Fig. 6.**
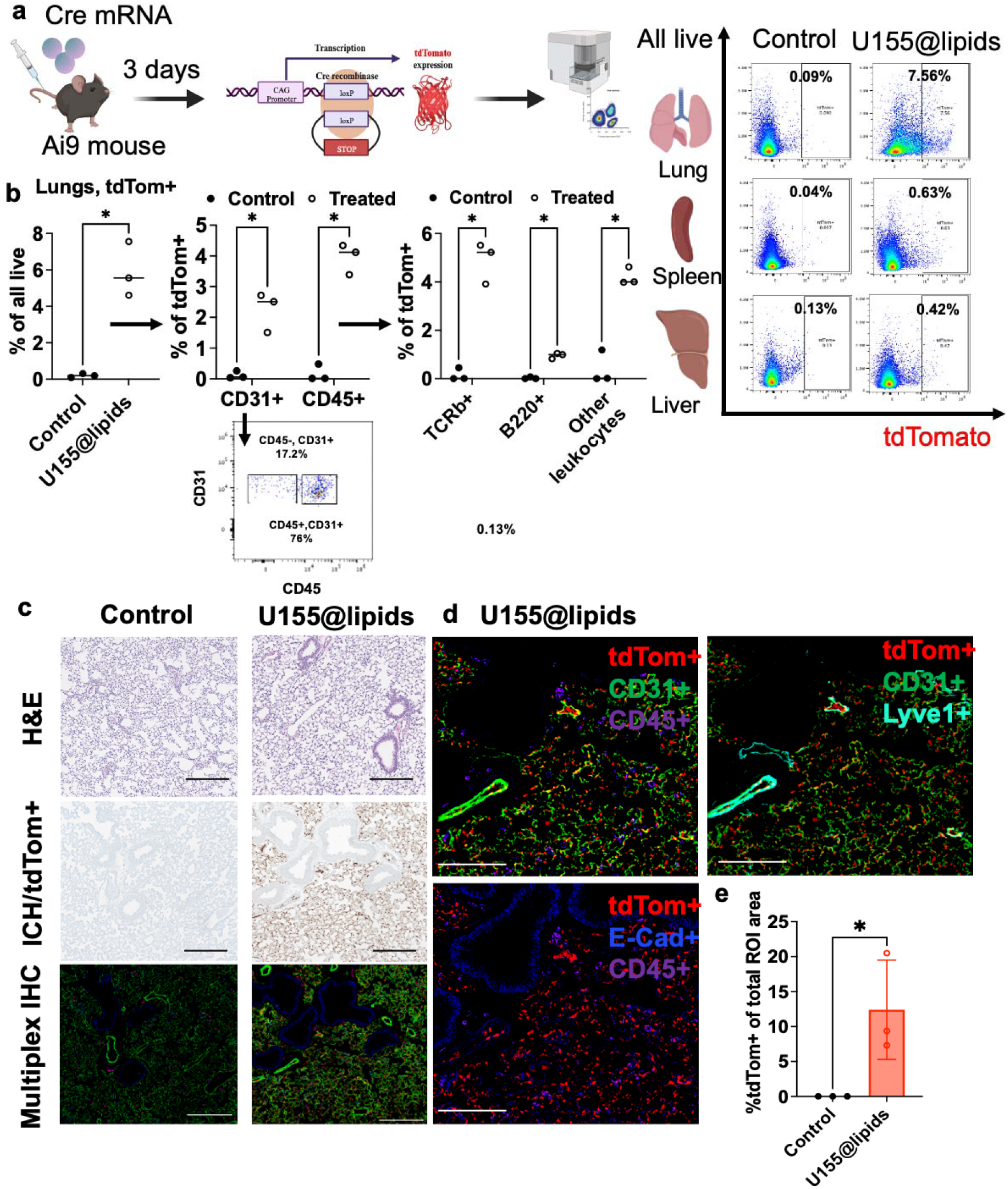
Assessment of *in vivo* Cre mRNA transfection. **a)** experimental scheme and represent tive tdTomato+ cells as a percentage of the total live cell population in each of several major organs: Ai9 mice were injected with U155@lipids encapsulating Cre mRNA, and tdTomato expression in single cell–level was quantified 3 days after injection using flow cytometry; **b)** quantification of tdTomato+ cells in the lungs in different cell types (tdTomato+ cells as a percentage of the overall population of each cell type). Data are presented as Means ± SD (n=3), (p<0.05); **c)** representative images of paraffin- embedded lung sections, which were stained with H&E and antibodies for multiplex IHC. Scale bar is 200 μm; **d)** Multiplex IHC of lung sections; **e)** quantification of IHC in **c**. Data are presented as Means ± SD (n=3 biological replicates). Representative image of how QuPath’s tool select tdTomato+ stained areas is in Supporting information Fig. S13.

### Treatment with IL-12 mRNA loaded hybrid U155@lipids confers a significant survival advantage in Lewis Lung cancer model

As U155@lipids transfects mostly lung endothelial and immune cells we endeavored to apply our nanoparticles for immunotherapeutic purposes in a mouse model of lungs cancer. Interleukin-12 (IL-12) is cytokine noted for its anti-tumor activity in a variety of preclinical models.^82^ Previously it was reported that IL-12 mRNA delivery may be an effective immunotherapy against multiple cancer types.^83–87^ Upon delivery, the host cells translate the IL-12 mRNA into IL-12 protein. IL-12 is a cytokine that plays a crucial role in polarizing the immune system towards Th1/cytotoxic immune milieu, associated with functional anti-tumor immune response. IL-12 is particularly necessary for polarizing T cells through which the tumor microenvironment can also be changed (Fig. 7a). To check the therapeutic potential of our platform we established an orthotropic lung cancer model by i.v. injection of Lewis lung carcinoma cells expressing luciferase (LLC-Lum) and evaluated treatment with IL-12 mRNA (Fig. 7b).

**Fig. 7.**
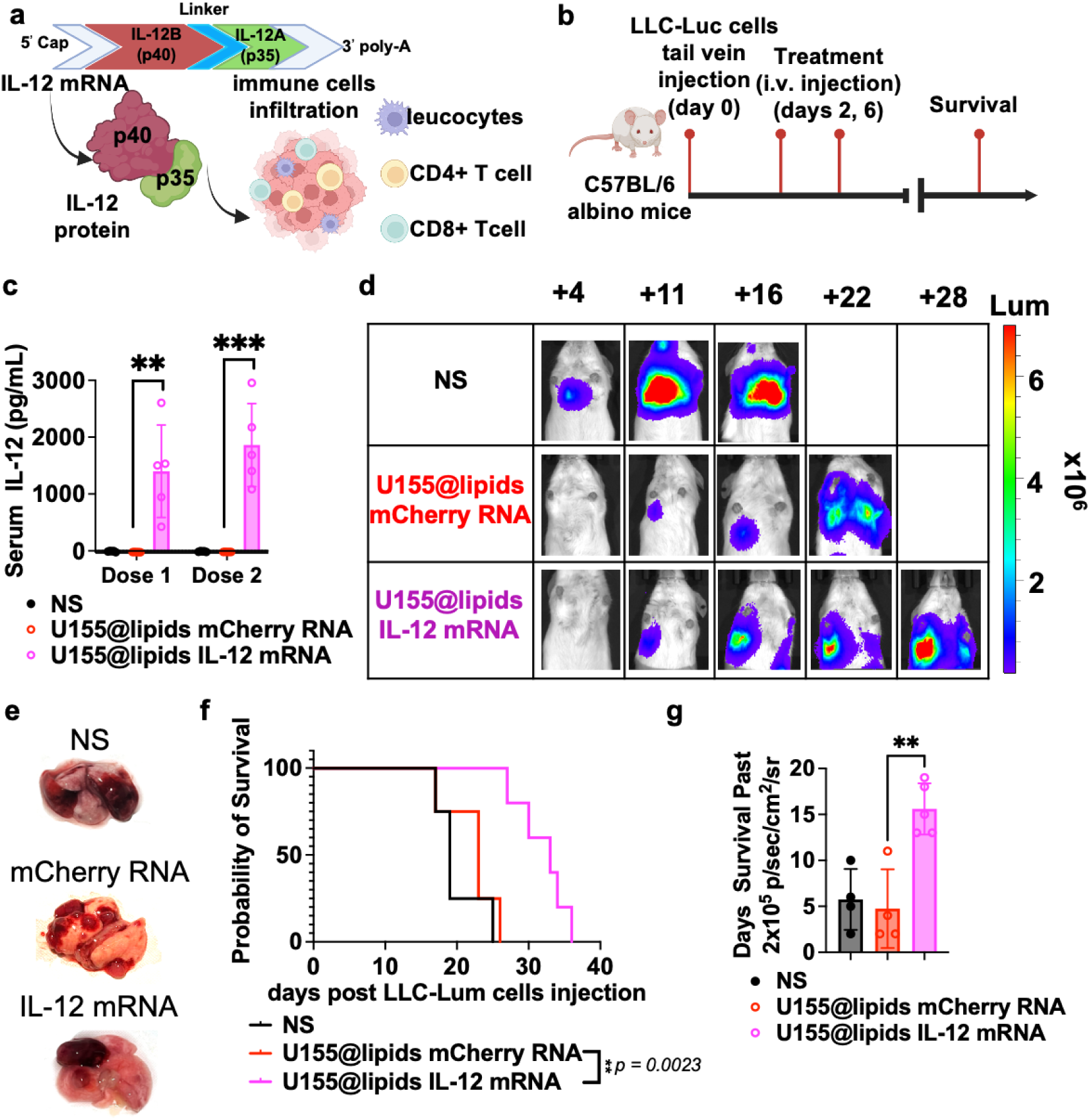
Assessment of *in vivo* IL-12 mRNA immunotherapy. **a)** structure of IL-12 mRNA and protein; **b)** experimental scheme of treatment; **c)** serum IL-12p70 concentration 24 h post 1^st^ and 2^nd^ i.v. injections of IL-12 mRNA and mCherry RNA (5 ug dose per mouse), normal saline (NS). Data are presented as Means ± SD (n=5), (**p<0.005, ***p<0.0005); **d)** Representative images of bioluminescence in lungs upon time; **e)** representative images of *ex vivo* lungs; **f)** Kaplan-Meier survival analysis comparing overall survival (OS) between control (NS, *n* = 4), mCherry RNA (U155@lipids mCherry RNA, n=4), and IL-12-mRNA treated (U155@lipids IL-12 mRNA, n = 5). P-values were calculated using log-rank test; **g)** days survival past in vivo bioluminescence signal reached 2×10^5^ **p<0.005. p/sec/cm^2^ /sr. n=4-5,

We explored IL-12 protein expression (Fig. 7c) in serum 24 h post i.v. dose administrations. Notably, animals treated with IL-12 mRNA exhibited substantial protein levels, in contrast to the negligible cytokine presence with PBS or mCherry RNA treatments (Fig. 7c). Interestingly, IL-12 cytokine concentration was approx. the same after the first and second doses, once again confirming the effectiveness of our platform for multiple dosage administrations. Tumor growth was monitored by bioluminescence signal (Fig.7d,e, supporting information Fig. S15). Administration (i.v.) of IL-12 mRNA loaded U155@lipids significantly delayed tumor progression prolonged overall survival of mice compared to PBS and mCherry RNA loaded U155@lipids controls (Fig. 7f). While all mice eventually succumbed to tumor burden, mice receiving IL-12 survived significantly longer even once larger tumors were established (2×10^5^ p/sec/cm2/sr) consistent with the presence of an active anti-tumor immune response (Fig. 7g). Despite transient weight loss for the first days after IL-12 mRNA treatment, animals recovered and, unlike the control groups, maintained weight for at least 10 days post the second dose (supporting information Fig.S16). Notably two doses of mCherry RNA control also induced a delay in tumor progression and but did not significantly induce IL-12, extend overall survival, nor slow time to death post establishment of large tumors (Fig.7 c,d,f,g-f). Similar results were previously reported, thus overall^83^, our results further support that IL-12 initiated immune stimulation provided by RNA delivery, could yield a significant yet transient therapeutic benefit.

### Delivering of CFTR mRNA restores CFTR-mediated chloride efflux in CFTR KO mice

To assess our level of success in developing a broadly employable therapeutic platform across mRNA sizes, next, we sought to deliver a therapeutic mRNA larger than Fluc (1922 b) or IL-12 (1617 b) mRNA and subsequently check the functionality of expressed protein. Delivery of CFTR mRNA (6132 b) is a potential therapeutic approach for cystic fibrosis, a genetic disorder characterized by the non-functioning of the CFTR protein. CFTR mRNA delivery aims to introduce functional mRNA into cells, allowing them to produce the CFTR protein (Fig. 8a). Based upon the latest updates from cystic fibrosis foundation, targeting multiple cell types for gene therapies including respiratory, intestinal and nasal epithelial cells, immune cells and stem cells, with CFTR mRNA treatment can is predicted to offer comprehensive therapeutic benefits and improve the quality of life for individuals with cystic fibrosis through restoration of a variety of functional activities in these varied cell types.^88,89^

**Fig. 8.**
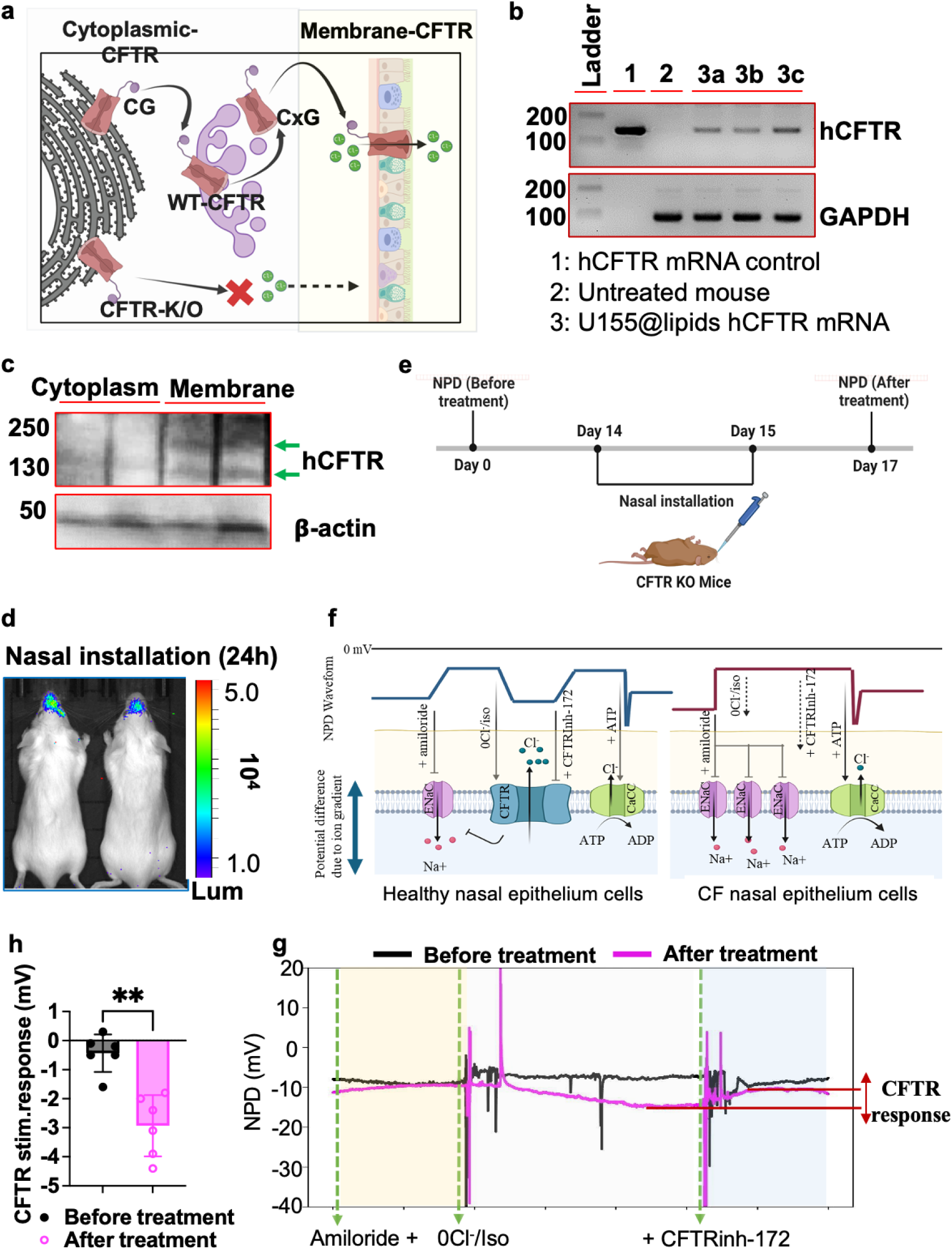
**a)** CFTR-WT is core-glycosylated (CG) in the endoplasmic reticulum, after which it moves through the trans-Golgi network to become complex-glycosylated (CxG) and reaches the plasma membrane, where it acts as a chloride channel; **b)** RT-PCR analysis of BALB/c mice lungs 4 h after i.v. injection of U155 lipids hCFTR mRNA. Negative control, untreated mouse; positive control, hCFTR mRNA; loading control, GAPDH; **c)** Western blot detection of hCFTR protein in untreated and treated CFTR KO mice lungs lysates. β-Actin was used as a loading control; **d)** bioluminescence signal 24 h post intranasal instillat on of U155@lipids Fluc mRNA (4 µg Fluc mRNA per mouse) in BALB/c mice, measured using IVIS experimental scheme of treatment **e)**; experimental scheme of treatment; **f)** schematic diagram illustrating the correlation between NPD traces and ion transport; **g)** representative NPD traces for a single CFTR KO mouse before and 48 hr post treatment with U155@lipids hCFTR mRNA (2 days nasal installation with total 8 µg mRNA dose per mouse); **h)** hCFTR response before and after treatment. Data are presented as Mean ± SD; **p < 0.005,

To test the utility of our platform to provide CFTR we, first, we injected via tail vein hCFTR mRNA (8 µg per mouse) loaded in our platform and 4 h later harvested lungs, extracted total RNA and generated cDNA libraries through RT-PCR. To prove the delivery of hCFTR mRNA, PCR amplification of exon 11 of human CFTR was done as described previously^90^ (Fig. 8b) demonstrating clear expression of hCFTR in lung. The CFTR protein undergoes post-translational modifications, becomes core-glycosylated (135 kDa) in the endoplasmic reticulum and is further modified to a functional complex through extensive glycosylation (180 kDa). To be functional it next needs to move through the trans-Golgi network before reaching its functional destination at the plasma membrane (Fig. 8a).^91^ This folding process is crucial for CFTR protein proper function as a chloride ion channel and partly depends on integrity of delivered mRNA. Using western blot analysis, we confirmed efficient delivery of mRNA and expression of functional CFTR protein in the lungs after treatment in CFTR knockout (KO) mice. We compared the lysates from cytoplasm and membrane and the presence of CFTR enrichment in membrane strongly indicates that exogenous hCFTR mRNA is translated and undergoes post-translational modification and subsequent membrane association (Fig. 8c).

By expanding our route of administration to include intranasal administration of nanoparticles we demonstrated significant transfection of nasal epithelium post nasal installation (Fig. 8d). To evaluate the to evaluate *in vivo* functionality of expressed CFTR protein we measured nasal potential difference (NPD) in CFTR KO mice before and after nasal instillation treatment (Fig. 8e). The difference in electrical potential across conductive epithelium arises from the movement of ions across the epithelial layer and is mainly influenced by the activity of three key channels: the epithelial sodium channel (ENaC), CFTR, and calcium-activated chloride channels (Fig. 8f).^92^ In CFTR KO mice, there is inadequate transport of chloride ions, leading to increased absorption of sodium and an exaggerated response to substances like amiloride, which inhibit ENaC. Notably, inhibition of CFTR with CFTRInh-172 does not produce any observable effects in CFTR KO mice due to the absence of the CFTR protein (Fig. 8g,h). Once we delivered hCFTR mRNA (total 8 µg per mouse) mice exhibited polarization in response to CFTR stimulation (Fig. 8g), followed by inhibition after addition of CFTRInh-172, confirming that the observed response was CFTR-dependent (Fig. 8h). Collectively, we demonstrated that our platform successfully delivered the large hCFTR mRNA to the lungs preserving functionality of the cargo and translated protein.

### Hybrid U155@lipids enable CRISPR-Cas9 editing in the lungs

Demonstrating a high lungs tropism upon i.v. delivery and achieving successful transfection in endothelial and T cells with *in vivo* tolerability, we proceeded to investigate the potential of employing U155@lipids for efficient delivery of CRISPR-Cas9 systems to the lungs in the Ai9 mouse strain. To do this we delivered Cas9 and sgRNAs targeting the stop cassettes present in the ROSA26 locus, impeding expression of tdTomato expression. Introduction of insertions or deletions in tdTomato STOP cassettes (Fig. 9a) results in a frame shift which negates the STOP codon and results in tdTomato protein expression. We injected via tail vein U155@lipids encapsulated Cas9 mRNA+sgRNA (1/1 w/w) at dose of 19 µg total RNA and quantified gene editing on day 9 post injection (Fig.9a). As before multiplex IHC of lung tissue revealed editing only in CD31+ endothelial cells (Fig. 6b). Notably Cre-recombinase editing was at least 15-fold more efficient than the CRISPR-Cas9 system (Fig. 6d and Fig. 9c), concentration of potentially edited CD45+ cells detected by IHC was expected to be lower. To prove that tdTomato expression was caused by indels of three STOP codons, we used the next-generation sequencing analysis of DNA, extracted from lungs. Editing was quantified as sum of the insertions and deletions, normalized to the number of reads, resulting in a % of reads containing indels. We found that treated mice displayed 5.6 ± 2.4 % of editing (Fig. 9d).

**Fig. 9.**
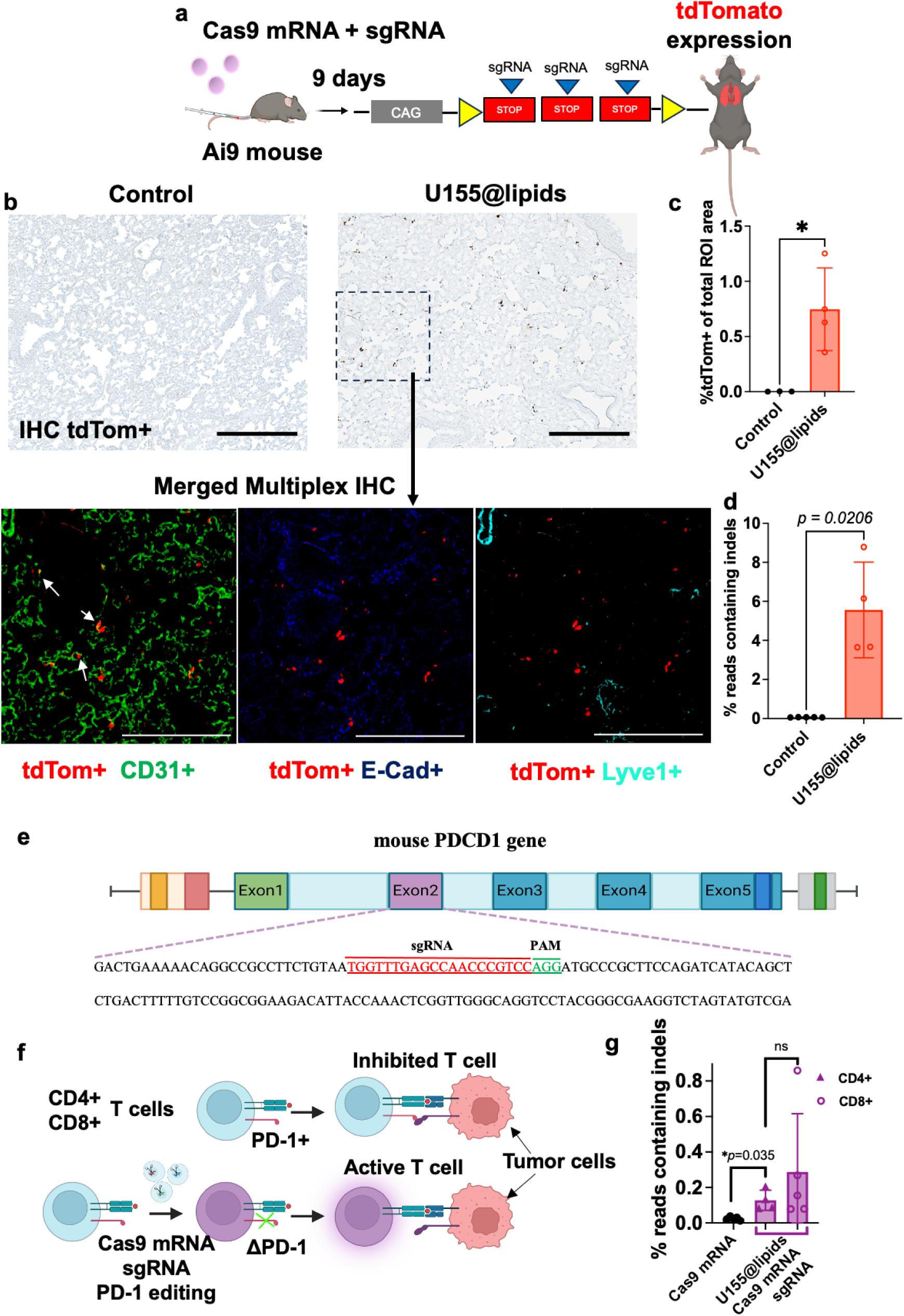
Assessment of *in vivo* CRISPR-Cas9 editing. **a)** experimental scheme: Ai9 mice were injected with U155@lipids encapsulating Cas9 mRNA/sgRNA (1/1 wt/wt ratio), and 9 days after injection lungs were collected and analyzed with next-generation sequencing (NGS) and IHC; **b)** representative images of paraffin-embedded lung sections, which were stained with antibodies for multiplex IHC. White arrows point out the tdTomato and CD31+ co-localization. Scale bar is 200 μm; **c)** quantification of IHC in **b**. Data are presented as Means ± SD (n=3-4 biological replicates). Representative image of how QuPath’s tool select tdTomato+ stained areas is in Supplementary Figure S12; **d)** quantification of editing events in lungs by NGS. ata are presented as Means ± SD (n=4-5 biological replicates); **e**) schematic of the Cas9 target site within the mouse PDCD-1 gene at the exon2 location; **f)** schematic of interaction of PD-1 positive and ΔPD-1 knockout T cells with tumor cells; **g)** quantific tion of editing events in CD4+/CD8+ T cells by NGS. Data are presented as Means ± SD (n=4-5 biological replicates).

Encourage by these results, we sought to extend the application of our particles to show functional editing in T cells. Administration of antibodies targeting PD-1 have significantly improved outcomes for cancer patients through removal of the “checkpoint” signal initiated through PD1 expressed by T cells that normal limits their anti-tumor activities (Fig. 9f). Ablation of PD1 signalling has been demonstrated to enhance not only the activity of resident T cells but also increase the persistence of transferred CAR-T cells.^93,94^ Thus, we sought to design a guide which provides deletion and frame shift in mouse exon 2 of the PD1 locus, permanently blocking future expression of the PD-1 protein (Fig. 9e). I.v. administration of one dose (10 µg total RNA, Cas9 mRNA/sgRNA w/w ratio of 1/1) of U155@lipids nanoparticles after pre-activation with a single i.v. dose of Il-12 mRNA (5 µg) showed up to 0.29% of gene editing in both CD4+/CD8+ T cells based on NGS analysis (Fig. 9g). Notably, to our knowledge this is the first report of editing of the PD-1 gene *in vivo* through a non-viral delivery of Cas9 mRNA system. These results open more possibilities of application of our platform.

## Discussion

Non-viral systems for mRNA delivery to the lungs have gained significant interest due to their potential in treating various respiratory diseases, including cystic fibrosis, lung cancer, and infectious diseases like COVID-19. Nowadays dozens of lipids and polymer-based platforms with lung tropism are designed for various routes of administration, however, most of them suffer from side effects such as immunogenicity and lung embolism due to highly positive charge. Addressing these challenges is crucial for the successful development of safe and effective mRNA delivery systems for lung-targeted therapies.

We have successfully implemented a highly efficient method for synthesizing cationic PEI-based polymers utilizing the split-Ugi reaction. By formulating a structurally diverse library of these polymers into polyplexes and screening them *in vitro* and *in vivo*, we have resolved the problem of balancing particles stability with cargo release, which are shown to depend on the degree of polymerization and polymer hydrophobicity. In particular, the incorporation of lipid-like moieties at high modification degrees (∼50% of polymer repeat units) was identified to yield lipopolymers with high transfection performance. Additionally, the introduced tertiary amine units of the polymer may be beneficial as they have lower proton buffering capacity at physiological pH compared to primary and secondary amines, resulting in reduced osmotic stress and decreased cellular toxicity.^95,96^ In future work, the prospect of synthesizing these split-Ugi derived polymers with biodegradable backbone units seems promising.

It is worth noting, while clearly proven effective, both lipopolymer structure and lipid composition of the final formulation can be further improved but the main goal of the work was to demonstrate the possible applications and perspectives of the offered approaches in mRNA delivery platform formulating.

Using a two-step approach, we produced lipopolymer-lipid hybrid nanoparticles. These nanoparticles selectively deliver and induce effective mRNA expression in lung endothelium and immune cells, including T and B cells, with minimal *in vivo* toxicity after i.v. administration. Remarkably, U155@lipids demonstrate nearly 300-fold higher potency in systemic mRNA delivery to the lungs compared to *in vivo*-JetPEI^®^. We showed that U155@lipids accumulation in the lungs could potentially be elucidated as ‘passive’ organ-selective delivery. Similar to previous reports, pre-injection of blank nanoparticles decreases the liver accumulation, improving U155@lipids mRNA delivery to the lungs. Notably, our platform demonstrated efficient multiple dosages administration which is useful in the case of gene editing with complicated processes required high doses for therapeutic effect.

Another advantage of our platform and formulation approach is the delivery of therapeutic mRNA of various size for effective expression of functional proteins. Thus, systemic delivery of IL-12 mRNA loaded U155@lipids significantly delayed lungs tumor progression and prolonged overall survival of mice. We expect that this therapy might be improved by combination with other strategies including PD-1 gene knockout, which also was demonstrated using our platform. However, directly knocking out the PD-1 gene in T cells may have significant consequences. While it might enhance the immune response against cancer, it could also lead to autoimmunity, where the immune system attacks healthy cells and tissues in the body. Therefore, this approach and further application of our platform need to be carefully studied and potentially combined with other strategies to minimize potential autoimmune side effects.

The capacity to deliver mRNA to the lungs through various routes of administration also increases the number of cell types that can be affected, enabling also treatment of cystic fibrosis. Our platform successfully delivered large hCFTR mRNA to the lungs preserving functionality of the cargo and translated protein upon i.v. administration. Also, instead of i.v. injection, intratracheal installation could be implemented (supporting information Fig. S17). However, for further improvement of therapeutic effect, the CFTR mRNA sequence could be optimized.^97^ Furthermore, U155@lipids demonstrated the efficient delivery of CRISPR-Cas9 system to the lungs, resulting in significant gene editing within tissues. This could potentially enable a more efficient strategy for cystic fibrosis treatment.

To conclude, our findings underscore the tremendous potential of our synthetic and formulating approaches for mRNA delivery and gene editing in the lungs, holding promise for a wide range of therapeutic applications.

## Materials and methods

Octylamine, decylamine, dodecylamine, cyclohexanecarboxaldehyde, octanal, decanal, benzylamine and trans,cis-2,6-nonadienal were obtained from TCI Chemicals and used as received. Ethyl formate, hexanal, dodecanal, formaldehyde (37 % in H_2_O), acetic acid, hexanoic acid, 3-(dimethylamino)propionic acid hydrochloride, cyclohexylisocyanide, ethyl isocyanoacetate and 2-morpholinoethyl isocyanide, cholesterol, 30% Hydrogen peroxide solution (7722-84-1), citric buffer (10x) antigen retriever were obtained from Sigma Aldrich and used as received. L-α-phosphatidylcholine (Soy-PC), 1,2-distearoyl-sn-glycero-3-phospho-(1’-rac-glycerol) (sodium salt) (DSPG) were purchased from AvantiPolarLipids. 1,2-Dimyristoyl-rac-glycero-3-methylpolyoxyethylene (DMG-PEG2000) was obtained from NOF American Corporation. DiD’ solid (1,1’-Dioctadecyl-3,3,3’,3’-Tetramethylindodicarbocyanine, 4-Chlorobenzenesulfonate Salt) was purchased. From ThermoFisher Scientific. AMEC Red Substrate Kit, Peroxidase (HRP) (SK-4285) was obtained from Vector Laboratories. CleanCap Fluc mRNA, mCherry mRNA, Cre mRNA, customized IL-12 mRNA, human CFTR mRNA (NCBI: NM_000492.3, custom-made), Cas9 mRNA, each with fully substituted uridine by pseudouridine and cytidine by 5-methylcytidine, were purchased from TriLink Biotechnologies. Guide RNA for tdTomato and PD-1 gene editing, all primers for PCR were ordered from Integrated DNA Technologies.

Acetonitrile, benzonitrile, benzylbromide, methyl triflate and 2-ethyl-2-oxazoline were obtained from Sigma Aldrich and dried over CaH, then purified by distillation before use. **NMR** spectra were recorded on a Bruker Ultrashield 500 MHz Plus system at 25 °C using deuterated solvents obtained from Sigma-Aldrich. **MALDI-ToF-MS** was performed on a Shimadzu Axima Performance instrument in or positive-reflector mode. *Trans*-2-[3-(4-*tert*-Butylphenyl)-2-methyl-2- propenylidene]malononitrile (DCTB) (100 mg mL^-1^ in ACN) was used as the matrix without further purification (Sigma-Aldrich). NaTFA salt was used as the ionization agent (1 mg mL^-1^ in MeOH). Matrix, polymer, and salt solutions were mixed in a 1:1:0.5 volume ratio and then 1 μL of the mixture was deposited onto a ground steel target plate before insertion into the ion source chamber. The instrument was calibrated against a poly(ethylene glycol) methyl ether standard (Sigma Aldrich, Average Mn = 2,000 g mol^-1^) prepared under the same conditions with DCTB matrix.

### SEC-RI chromatography

Poly(ethyloxazoline) samples were analysed with the Agilent 1260 Infinity II chromatography system with Stryagels HR2, HR4 and HT5 columns and Agilent 1260 infinity RI detector was used for the SEC studies. DMF + 0.1% LiBr was used as an eluent. The flow rate was 0.8 mL/min. The column was thermostated at 40 C. The Ugi modified polymers were instead analysed on a system using an eluent mixture of CHCl3, IPA and triethylamine (TEA) in ratios of 94:4:2. The system consisted of Waters 515 HPLC pump, Biotech DEGASi GPC Degasser, Waters 717 plus Autosampler and Waters 2410 Differential Refractometer together with Waters Styragel HR 2, HT 3, and HT 4 7,8x300 mm and guard column. The flow rate was set to 0.800 ml/min and the columns thermostated at 30 C PMMA standards obtained from Polymer Standards Service were used for calibration.

### Animals

Female BALB/C, male C57BL/6 albino (Charles River Laboratory) and female CFTR-/- tm1Unc Tg(FABPCFTR)1 Jaw/J double-transgenic CFTR KO, male B6.Cg-Gt(ROSA)26Sortm9(CAG-tdTomato)Hze/J (Ai9) (The Jackson Laboratory) mice, aged 8 weeks, were used in the study. All animal care and experimental procedures were approved by the Institutional Animal Care and Use Committee (IACUC, protocol # IP00001707, #IP00002318, # IP00002286) of Oregon Health and Sciences University.

### Synthesis of benzylisocyanide

Selected isocyanides were synthesized using a modified literature procedure.^98^ Benzylamine (5 g, 46.7 mmol) was dissolved in ethyl formate (11.3 ml, 140 mmol) in a round bottom flask and refluxed for 3 hours at an oil bath temperature of 65 C. The reaction mixture was concentrated under reduced pressure.

The formate intermediate was dissolved in dichloromethane (DCM) (30 mL) under nitrogen atmosphere, triethylamine (32.5 mL, 233 mmol) was added, cooled with an ice water bath followed by dropwise addition of a solution of phosphorus oxychloride (4.56 mL, 49 mmol) in DCM (5 mL). The reaction mixture was stirred for 2 hours and allowed to warm to room temperature, and then purified directly by column chromatography over silica. Diethyl ether was initially used as the mobile phase, switching to a 50 % DCM/Et_2_O mixture after elution of approximately one column volume. The product fractions were combined, concentrated and then purified by distillation under vacuum to yield a pale-yellow oil (yield = 2.3 g, 42 %). ^1^H NMR (500 MHz, CDCl_3_) δ 7.41 (m, 5H, C_5_<u>H</u>_5_), 4.67 (2H, s, C<u>H</u>_2_).

### Synthesis of alkylisocyanides

The same protocol described above for benzylisocyanide was employed using 5 g of the alkylamine reagent, modifying quantities of the other reagents accordingly. For column chromatography a hexane / ethyl acetate (9:1 to 4:1) mixture was used. Only octylisocyanide was further purified by distillation.

**Octylisocyanide.** Yield = 3 g, 56 %. Distilled at 60 C, 0.2 mbar. ^1^H NMR (500 MHz, CDCl_3_) δ 3.40 (tt, 2H, J = 1.9, 6.7 Hz, C<u>H</u>_2_NC), 1.70 (2H, m, C<u>H</u>_2_CH_2_NC), 1.45 (2H, m, C<u>H</u>_2_(CH_2_)_2_NC) 1.32 (8H, m, CH_3_(C<u>H</u>_2_)_4_) 0.91 (3H, t, J = 7.0 Hz, C<u>H</u>_3_).

**Decylisoycanide** Yield = 4.1g, 77 %. ^1^H NMR (500 MHz, CDCl_3_) δ 3.40 (tt, 2H, J = 1.9, 6.7 Hz, C<u>H</u>_2_NC), 1.70 (2H, m, C<u>H</u>_2_CH_2_NC), 1.45 (2H, m, C<u>H</u>_2_(CH_2_)_2_NC) 1.32 (8H, m, CH_3_(C<u>H</u>_2_)_4_) 0.91 (3H, t, J = 7.0 Hz, C<u>H</u>_3_).

**Dodecylisocyanide** Yield = 3.8 g, 72 %. ^1^H NMR (500 MHz, CDCl_3_) δ 3.40 (m, 2H, C<u>H</u>_2_NC), 1.70 (2H, m, C<u>H</u>_2_CH_2_NC), 1.46 (2H, m, C<u>H</u>_2_(CH_2_)_2_NC) 1.32 (16H, m, CH_3_(C<u>H</u>_2_)_8_) 0.91 (3H, t, J = 6.9 Hz, C<u>H</u>_3_).

^1^H NMR are presented in Supporting information Fig. S18-S21.

### Synthesis of poly(ethylene imine)s

#### Cationic ring opening polymerization of 2-ethyl-2-oxazoline

2-Ethyl-2-oxazoline (15 g, 151.3 mmol), benzonitrile (35.1 mL) and methyl triflate (0.69 – 4.32 mmol) were added to a dry Schlenk flask under inert atmosphere, and then stirred at 80 C for 4 hours. Molar equivalents of methyl triflate (MeOTf) were altered to target degrees of polymerization 35, 85 and 220 accordingly. The reaction mixture was cooled to 40 C, terminated by addition of benzylamine (10 eq. with respect to MeOTf) and left to stir overnight. The polymer was precipitated three times from diethyl ether and dried under vacuum to yield a colorless powder.

For the preparation of the lowest degree of polymerization PEtOx_15_, the initiator benzyl bromide was used in acetonitrile and the termination carried out with 1M KOH solution instead. Yield = 4.9 g (88 %). SEC and MALD-ToF results are presented in Supporting information Fig. S22 and Table S2.

### Hydrolysis of poly(2-ethyl-2-oxazoline)

Poly(2-ethly-2-oxazoline) was dissolved in 3M HCl (75 mL) in a round bottom flask fitted with a stir bar and refluxed overnight for 18 hours. The mixture was cooled to room temperature, then adjusted to pH 10 by addition of 4 M NaOH causing precipitation of the polymer. The solid was collected by centrifugation and washed five times with distilled water collecting again by centrifugation between washes. The solid was dried under vacuum to yield a colorless powder. ^1^H NMR is presented in supporting information Fig. S23.

### Ugi modification of poly(ethylenimine)

Reactions were run targeting total modification amounts of PEI units of 25-100%, assuming the reaction forms the split-Ugi product whereby 2 molar equivalents of PEI secondary amines are required with respect to the other reagents. The quantities of reagent were calculated as follows taking the repeat unit of PEI as 1 eq: 25% target modification: 0.125 eq. aldehyde, 0.125 eq. isocyanide, 1 eq. carboxylic acid. 50 % target modification: 0.25 eq. aldehyde, 0.25 eq. isocyanide, 1 eq. carboxylic acid. 100 % target modification: 0.5 eq. aldehyde, 0.5 eq. isocyanide, 1 eq. carboxylic acid.

PEI was dissolved in EtOH/H_2_O (9/1) at a concentration of 50 g/L polymer, and stirred with heating at 50 C. When targeting the highest modification density, the concentration was lowered to 20 g/L polymer to ensure sufficient solubility. The aldehyde reagent was added to the PEI solution and stirred for 30 minutes, followed by addition of the isocyanide and then the acid. The mixture was stirred overnight with continued heating of 50 C, then cooled to room temperature, diluted with ethanol, and transferred to a dialysis membrane (regenerated cellulose, MWCO = 1 kDa). Dialysis was performed first once against ethanol, and then twice against deionized water exchanging solvents every 24 h. The solution was freeze dried to yield the product, typically as a pale orange solid / oil. Alternatively, in early batches of samples (U1 – U11) the polymers were purified by precipitating three times into ice cold diethyl ether, resolubilizing in ethanol between steps. This procedure was unreliable, however, for the more hydrophobic derivatives which showed some degree of solubility in diethyl ether or hexane and thus dialysis was selected as the purification method of choice. NMR spectra are presented in supporting information Fig. S24-S30, Table S3.

### Polyplex preparation

For *in vitro* and *in vivo* screening polyplexes were formulated from the synthesized PEI- derivatives library via ethanol injection method. Briefly, a solution of the polymer in ethanol (2 g/L) was combined with an aqueous phase containing mRNA (28.6 g/L) in an acetate buffer (25 mM, pH 5, IS 154 mM) at the ratio water to ethanol 7/1 (v/v), resulting in a 10/1 (w/w) polymer to mRNA ratio. The mixture was then left to incubate for 30 minutes, allowing the polyplexes to form. Finally, the polyplexes were neutralized using Tris-HCl buffer (25 mM, pH 7.4, IS 154 mM). For *in vivo* studies polyplexes were dialyzed against Tris-HCl buffer (25 mM, pH 7.4) for 2 hours at room temperature. Size distribution and polydispersity indexes (PDI) of polyplexes were determined with dynamic light scattering using Stunner (Unchained Labs, US) or Zetasizer Nano ZSP (Malvern Instruments, UK). Concentration of RNA in final formulation was assumed to be 100% yield.

### Hybrid polymer-lipid particles preparation

Hybrid polymer-lipid nanoparticles were formulated in two steps. Before formulating, all lipids with specified molar ratios were dissolved and mixed in ethanol to form a complete lipid mix solution. Separately, polymer was dissolved in ethanol and mRNA was diluted in 25 mM acetate buffer (pH 5.0). Then, the ethanol polymer solution was rapidly mixed by vortexing with the aqueous buffer solution containing Fluc mRNA at a ratio of 20/1 (aqueous/ethanol, v/v) to achieve a final weight ratio of 10/1 (total polymer/mRNA, w/w). The resulting mixture was then left to incubate for 30 minutes, allowing the polyplexes to form. Next, the aqueous solution of polyplexes was combined with ethanol phase, containing DSPG, soy PC, cholesterol and DMG-PEG2000 lipids in molar ratio 22/23/50/5, by microfluidic mixing using NanoAssemblr Ignite+ (Precision Nanosystems). Polymer to lipids ratio was 2/1/(w/w). Final hybrid particles were dialyzed 3-4 hours against Tris-HCl buffer (25 mM, pH 7.4) and concentrated with 10-kDa Amicon Ultra centrifuge filters (Millipore, Burlington, MA). Size distribution and PDI of polyplexes were determined with dynamic light scattering using Stunner (Unchained Labs, US) or Zetasizer Nano ZSP (Malvern Instruments, UK). RNA concentration in final formulation was measured with RiboGreen kit according to the manufacturer’s protocol with some modifications. Samples were diluted to 100 µg/L in 1× TE buffer with 2 g/L heparin (Sigma-Aldrich, U.K.). Standard solutions from the corresponding RNA stock were also prepared in a 1× TE buffer with 2 g/L heparin to account for any variation in fluorescence. Ribogreen reagent was diluted 2000-fold in 1× TE buffer. RNA encapsulation of samples was determined by comparing the signal of the RNA-binding fluorescent dye RiboGreen in the absence and presence of a detergent (2 % Triton X- 100). All samples were incubated 15 min at 37 °C before addition of reagent. In the absence of detergent, the signal comes only from unencapsulated RNA. In the presence of detergent, the particles were disrupted so that the measured signal comes from the total. Fluorescence was measured at λ_ex_ = 485 nm and λ_em_ = 530 nm.

To evaluate particles *in vivo* biodistribution, DiD dye was added to lipids mixture at 0.1% molar of total lipids concentration. For pretreatment studies, blank nanoparticles were prepared according to the protocol without mRNA in acetate buffer.

### Cryo-transmission electron microscopy (TEM)

Cryo-TEM images were captured with Falcon III and K3 Summit cameras with DED at 300 kV. The Vitrobot Mark IV system (FEI) was used to plunge-freeze a copper lacey carbon film-coated grid (Quantifoil, R1.2/1.3 300 Cu mesh). U155@lipids (10 µL) was dispensed onto the glow discharged grids in the Vitrobot chamber maintained at a temperature of 23 °C and a relative humidity of 100% to freeze the samples. The sample was incubated for 30 seconds before being blotted with filter paper for 3 seconds before being submerged in liquid ethane cooled by liquid nitrogen. The frozen grids were clipped. The images were taken at an electron dose of 15-20 e − /Å^2^ using 45,000 nominal magnifications with 1.5 binning then processed and analyzed using ImageJ (Fiji ImageJ2 version: 2.9.0/1.53t).

### TNS assay

TNS assay was performed as described previously.^99^ The McIlvaine citric-phosphate buffer was used to prepare buffers at various pH values between about 3 and 9 for determining apparent pKa. Then a stock of 300 μM 6-(p-toluidino)-2-naphthalenesulfonic acid sodium salt (TNS reagent) in DMSO was prepared. Buffer solution (90 μL) of was added to wells. Then 3.3 μL of U155 polyplexes or U155@lipids sample and 2 μL of 300 μM TNS reagent solution were added. Each well was then carefully mixed, and fluorescence was measured. With the resulting fluorescence values, a sigmoidal plot of fluorescence versus buffer pH was created.

### *In vivo*-JetPEI^®^ formulation

Complexes of mRNA with *In vivo*-JetPEI^®^ (Polyplus-transfection) were prepared according to the manufacturer’s instructions. Fluc mRNA (40 µg) and 6.4 µl *In vivo*-JetPEI^®^ (N/P = 8) were each diluted in 200 µL of 5% sterile D-glucose. Both solutions were mixed, followed by a 10-min incubation at room temperature.

### *In vitro* transfection efficacy screening

HeLa cells were plated in white, clear-bottom 384-well plates (2000 cells per well in 50 µL of complete DMEM media) and allowed to adhere overnight. Then, cells were treated with polyplexes loaded with FLuc mRNA (100 ng per well). Cell viability results (CellTiter-Fluor, Promega), and luciferase expression data (ONE-Glo Luciferase Assay, Promega) were collected 24 h post-treatment using a microplate reader. Luminescent readout (in relative luminescence units) was normalized by cell counts commensurate with fluorescence (relative fluorescence units, RFU).

### Endosomal escape studies

HEK293T/17 Gal9-GFP reporter cells were seeded (10,000 cells per well) in complete DMEM media in 96-wells black with clear-bottom plate. After overnight incubation, polyplexes loaded with mCherry RNA were added at a dose of 100 ng mRNA per well and incubated for 24 h. After incubation, media was gently aspirated, cells were then washed twice with PBS and fixed with 4% paraformaldehyde in PBS for 10 min at room temperature. Once cells were fixed, wells were gently washed with PBS two more times and DAPI (Thermo Fisher, Federal Way, WA) was then added at 1:1000 in PBS for nuclear staining. Following DAPI staining, cells were washed again two times and left in PBS. Reporter cells were imaged for GFP-positive puncta with a Fluorescent EVOS microscope with objective at 20× to report maximum intensity projections. Images were processed using ImageJ (Fiji ImageJ2 version: 2.9.0/1.53t).

### *In vivo* bioluminescence imaging

Fluc mRNA encapsulated in particles at a certain dose was injected *via* tail vein to female BALB/c mice. For bioluminescence imaging, mice received D-luciferin substrate (150 mg/kg) intraperitoneally and were imaged according to the manufacturer’s protocol. Image acquisition and analysis were performed using the IVIS Lumina XRMS and the manufacturer’s software (PerkinElmer). Mice were anesthetized by isoflurane inhalation during the procedure. For *ex-vivo* imaging, mice were sacrificed. Different organs were harvested, mocked in 1.5 g/L D-luciferin solution and analyzed.

### *In vivo* particles biodistribution

DiD-labeled nanoparticles were injected *via* tail vein to female BALB/c mice at the dose 5 µg of Fluc mRNA per animal. Mice were sacrificed 4 hr post-treatment, organs were harvested and imaged using the IVIS Lumina XRMS (λ_ex_/λ_em_ = 640 nm/670 nm). DiD concentration in formulation was measured in methanol and estimated using ε_640 nm_ = 2.45x10^5^ cm^-1^M^-1^. Control solution was prepared using DiD dye in DMSO, which was then diluted with DPBS to achieve final concentration of 1 % (v/v) DMSO.

### Lewis Lung cancer model treatment

For the Lewis Lung cancer model, LL/2-Luc2 cells (CRL-1642-LUC2, ATCC) in 100 µL PBS were tail vein injected into male C57BL/6 albino mice and randomly assigned to treatment and control groups. At days 2 and 6, IL-12 mRNA (5 µg), PBS (control) and mCherry mRNA (5 µg control) were i.v. injected. Tumor growth was monitored every 2-5 days by in vivo bioluminescence imaging. Animals were weighted every 2-5 days. For in vivo assessment of IL-12 mRNA transfection, mouse blood was collected 24 h post treatment and centrifuged at 10 000 xg for 15 min after 30 min of standing at room temperature. Concentration of IL-12 p70 in serum was determined using mouse ELISA Kit (Invitrogen) according to manufacturer’s instructions.

### NPD assay

For particle administration, mice were anesthetized with an intraperitoneal injection of a mixture of ketamine and xylazine (100 μg/10 μg/kg body weight). Particles were administered on two consecutive days to a single nostril (2 μL/application, 10 applications over 20 min, 4 µg /day).

NPD was measured using a modification of the previously described methods and using a previously described circuit.^92^ Following sedation with IP ketamine/xylazine, the mouse is orally intubated to protect the airway. Mice were positioned at a 15° head-down tilt, and a high-impedance voltmeter (World Precision Instruments, Sarasota, FL) attached to chloride pellet electrodes was used to measure the potential difference between an exploring nasal bridge and subcutaneous reference probe. A syringe pump continuously perfused solution into the nostril through a polyethylene tube stretched to approximately 0.5 mm in diameter (PE10, 0.28 mm inner diameter [ID]; Clay-Adams, BD, Sparks, MD). Solutions were perfused into the nasal cavity sequentially (Ringer’s, Amiloride, Gluconate substituted Ringer’s with isoproterenol, CFTR inhibitor, and 172Adenosine Triphosphate solution). DMSO (0.5%) was added to the CFTRInh-172 solution to improve solubility; this has been shown previously not to affect NPD measurements. NPD assay was performed prior to nanoparticle administration to record a baseline and then repeated 48 h after dosing.

### *In vivo* gene editing

For gene editing in mouse lungs with Cas9 mRNA/sgRNA in U155@lipids particles, Ai9 mice were injected with 100 μL of particles at dose total 19 μg per mouse (0.8 mg/kg; total RNA 1/1 mRNA/sgAi9, w/w). Animals were sacrificed on day 9 post-treatment, part of lungs was used for NGS analysis, another part of lungs was formalin-fixed, paraffin-embedded and analyzed with multiplex IHC In the sgRNA sequence, the asterisks (blue) and red font indicate the phosphorothioate bond and 2′-O-methyl ribonucleotides, respectively. A*A*G*UAAAACCUCUACAAAUGGUUUUAGAGCUAGAAAUAGCAAGUUAAAAU AAGGCUAGUCCGUUAUCAACUUGAAAAAGUGGCACCGAGUCGGUGCU*U*U*U

For PD-1 gene editing in mouse T cells with Cas9 mRNA/sgRNA, male C57BL/6 albino mice were treated (i.v.) with 5 μg dose of IL-12 mRNA loaded in U155@lipids and randomly assigned to treatment and control groups. At day 4, animals were treated (i.v.) with Cas9 mRNA/sgRNA system at dose total 10 μg per mouse (0.5 mg/kg; total RNA 1/1 mRNA/sgRNA, w/w) and Cas9 mRNA (control) at dose 5 μg per mouse. On day 10 post-treatment mice sacrificed, lungs and spleen were harvested, then dissociated and Cd4+/CD8+ T cells were isolated using EasySep^TM^ Mouse isolation kit (Stemcell Technologies) according to manufacturer’s instructions.

sgRNA sequence: TGGTTTGAGCCAACCCGTCC (exon 2, mouse).

### Flow cytometry studies

To study the biodistribution and cell type transfection by mRNA, Ai9 mice were intravenously injected with 10 µg Cre mRNA loaded into U155@lipids particles. On day 3 post-injection mice were sacrificed, lungs, spleen and liver were harvested and then dissociated. Single cells were generated with the organ dissociation protocol. Briefly, mouse organs were perfused with PBS and placed in a well of 12-wells plate with 0.5 ml of Click’s buffer on ice. Then 50 µl of collagenase IV and DNAse I mixture was added, and tissue was minced using tweezers and incubated for 30 min at 37 °C in 5% CO_2_ incubator, and then treated with 50 mM EDTA in D-PBS for another 5 min at 37 °C in 5% CO_2_ incubator. Subsequently, the digested tissue was filtered through a 70 µm nylon mesh strainer to collect single cells. Cells were washed using flow wash buffer (PBS containing 2 mM EDTA and 0.5% BSA) and collected by centrifugation at 450 x g for 5 min at 4 °C. The pelleted cells were resuspended in 100-200 µl of ACK lysis buffer (BioWhittaker, Lonza) for 5 min at room temperature and quenched with the addition of equal volume of 5 mM EDTA in PBS. After washing, cells were stained with live dead cell staining dye NIR and then incubated with Fc block (BioLegend). Cells were further stained with the following fluorochrome-conjugated antibodies: CD45-BUV395 (BD Horizon, #565967), B220-BUV805 (BD Horizon, #569199), CD31-BV605 (BD Horizon, #740356), TcrB-AF700 (BD Horizon, #560705). The stained cells were analyzed using Cytek Aurora. The gating strategy for flow cytometry analysis is shown in supporting information Fig. S11-S12.

### Multiplex Immunohistochemistry

For immunohistochemistry (IHC) studies, formalin-fixed, paraffin-embedded mouse lung samples were sectioned at 4 μm, deparaffinized and antigen retrieved with 10 mM citrate buffer for 20 min at 110 °C. Slides were then incubated 10 min in 3% peroxide in methanol, washed in TBS-tween buffer. Next, lungs sections were cyclically stained with antibodies in the following order: cycle 1 Rabbit RFP polyclonal antibodies (Biotium, 1:600 dilution, 1 h), cycle 2 Rabbit E-Cadherin (Cell Signaling, 24E10 #3195, dilution 1:200, 1 h), cycle 3 Rabbit CD31 (Abcam, ab182981, dilution 1:200, 1 h), cycle 4 Rabbit CD45 (Abcam, ab10558, dilution 1:200, 1 h), cycle 5 Rabbit Lyve1 (Abcam, ab33682, dilution 1:200, 1 h). Visualization was performed using the AMEC Red Substrate Kit, Peroxidase (HRP) (SK-4285) as instructed by the manufacturer. Digital scanning was performed using Aperio ImageScope AT2 (Leica Biosystems) at 20× magnification. Images were processed with ImageScopex64, QuPath and ImageJ (Fiji ImageJ2 version: 2.9.0/1.53t). Between each cycle tissue sections were treated with 95% ethanol for 10 min to wash blue protein staining and then 1% SDS, 25 mM glycine solution (pH 2-3) at 70 °C for 1 h.

### Hematoxylin and Eosin Staining (H&E)

Formalin-fixed, paraffin-embedded mouse lung and liver samples were sectioned at 4 μm. Lungs and liver tissues were first stained with hematoxylin for 30 seconds at room temperature and then rinsed in tap water. Then, tissue sections were decolorized in acid alcohol (10% acetic acid and 85% ethanol in water), followed by water washing. Next, sections were immersed in a saturated sodium bicarbonate solution, washed in water, and then immersed in 95% alcohol. Finally, staining was performed with Eosin-Phloxine for 10 seconds, followed by a dehydration step and cover slipping. Digital scanning was performed using Aperio ImageScope AT2 (Leica Biosystems) at 20× magnification. Images were processed with QuPath and ImageJ (Fiji ImageJ2 version: 2.9.0/1.53t).

### PAS-fast green staining

Formalin-fixed, paraffin-embedded mouse liver samples were sectioned at 4 μm, deparaffinized and hydrated to water. Then sections were oxidized in 0.5% periodic acid solution for 10 minutes and washed with water. Next, sections were placed in Schiff’s reagent for 30 min and wash in warm tap water for 5 minutes. To stain PAS-negative background elements tissue samples were incubate in Fast green (0.02% solution) for 30 seconds, following washing with water, dehydration with 95% and absolute alcohols, and cover-slipping. Digital scanning was performed using Aperio ImageScope AT2 (Leica Biosystems) at 20× magnification. Images were processed with QuPath and ImageJ (Fiji ImageJ2 version: 2.9.0/1.53t).

### Serum cytokines screening

Blood serum from female BALB/c mice 24 h post injection with 5 µg Fluc mRNA, loaded in U155@lipids, *via* tail vein injection was collected and analyzed by IDEXX BioAnalytics. Samples were tested on the Milliplex MAP Mouse Cytokine/Chemokine Magnetic Bead Panel (Millipore, Cat No. MCYTOMAG-70K-PMX) according to the kit protocol as qualified. Data were collected by xPONENT® 4.3 (Luminex) and data analysis was completed using BELYSA® 1.1.0 software. The data collected by the instrument software are expressed as Median Fluorescence Intensity (MFI). MFI values for each analyte are collected per each individual sample well. Analyte standards, quality controls, and sample MFI values were adjusted for background.

### PCR amplification and Western blot analysis

Lungs tissues were taken 4 h (for PCR analysis) and 48 h (for Western blot analysis) post hCFTR mRNA loaded particles administration (i.v.). For PCR analysis, RNA was isolated from lysates using the RNEasy Plus Micro Kit (QIAGEN), and cDNA libraries were generated by RT-PCR using the ProtoScript II cDNA Synthesis Kit (BioLabs) according to the manufacturers’ protocols. To demonstrate the presence of LNP-delivered hCFTR, primers were designed to amplify exon 11 (forward: 5’-AAC TGG AGC CTT CAG AGG GT-3’; reverse: 5’ -TTG GCA TGC TTT GAT GAC GC-3’) using GAPDH as a loading control (forward: 5’-AAC AGC AAC TCC CAC TCT TC -3’; reverse: 5’ – AGC CGT ATT CATTGT CAT ACC A -3’). PCR was carried out using KAPA kit (Invitrogen) in a 30-cycle reaction with a 10-min, 95 °C polymerase activation step. Each repeating cycle consisted of two steps: 15 s at 95 °C and 1 min at 60 °C. PCR products were analyzed using a 2% agarose gel. For Western blot analysis, tissue was homogenized and then lysed using Pierce^TM^ RIPA buffer (from cytoplasm extraction) and Tm-PER-100 (FIVEphoton Biochemicals) (from membrane extraction) containing HALT protease inhibitor (Thermo Scientific) at 4 °C. Total protein concentration was quantified by Micro BCA Protein Assay Kit according to the manufacturer’s instructions (Thermo Scientific). Samples and migration standards were prepared under reducing conditions and incubated for 10 min at 37 °C prior to loading. Wells were loaded at 400 ng (standards) or 100 µg total protein/well into a Novex WedgeWell 8% Bis-Tris gel (Invitrogen) and separated in Tris-glycine SDS running buffer (Novex). Transfer to a Nitrocellulose membrane (Novex) was performed in Tris-glycine transfer buffer (12 mM Tris-base, 100 mM glycine, 20% methanol) at 20 V for 90 min (Mini Blot Module, Invitrogen). For blocking, we used Super Block Blocking buffer (ThermoScientific). For staining Ms-anti-CFTR Ab596 (University of North Carolina), 1:2000, and Ms-anti-b-actin (Abcam), 1:10000 antibodies were used. The secondary antibody was horseradish peroxidase (HRP)-Gt-anti-Ms, 1:5000 (Abcam). For detection and imaging, SuperSignal West Pico Chemiluminescent Substrate (ThermoScientific) and a myECL Imager (model 62236X, Thermo Scientific) were used.

### Next generation sequencing (NGS)

NGS was performed via a two-step PCR as described previously.^100^ Genomic DNA (gDNA) was harvested from lung and spleen tissues and CD4+/CD8+ T cells using the Qiagen DNeasy Blood & Tissue kit (Qiagen, #69506) according to the manufacturer’s instructions. Briefly, genomic regions of interest were amplified using two primer sets specific to the first or second/third stop codon of the Ai9 loxp-stop-loxp cassette or to the exon 2 of the PD-1 gene of T cells (Table S4), and specific thermocycling conditions (Table S5). A second PCR was performed to add the necessary sequences to bind the amplicon to the Illumina flow cell (Table S5). The samples were then run on a gel, excised, pooled together, and harvested. The pooled library DNA concentration was quantified via qPCR using the KAPA Quantification Kit (Roche, #07960140001). After quantification the library was prepared for NGS using Illumina’s standard denature and dilute protocol. The NGS library was run on an Illumina MiniSeq using a 300-cycle mid output kit (Illumina, FC-420-1004). Editing was quantified using CRISPResso2 in standard mode, using the indel count in the “CRISPResso_quantification_of_editing_frequency.txt” file. The insertions and deletions were added together before being normalized to the number of reads, resulting in a % of reads containing indels.

### Statistical Analysis

All statistical analyses and significance were performed by GraphPad Prism 10 software (La Jolla, CA) or Microsoft Excel. An unpaired t-test was used to compare between two groups. Statistical significance among different groups and the data were considered statistically significant if p < 0.05.

### Display items

The images of nanoparticles, mice, syringes, organs (Figures 4a, 5a, 6a, 7a-b, 8, 9) were created with BioRender.com.

## Acknowledgments

Financial support from the Cystic Fibrosis Foundation (SAHAY19XX0), R01HL146736 from NHLBI (G.S), R01CA270783-01 (G.S) We thank PhD J. Kim, PhD Y. Eygeris, PhD A. Mukherjee for helpful advice and electron microscopy acquisition at Multiscale Microscopy Core (MMC), Oregon Health & Sciences University.

## Conflict of interest

K.Y. V, A.K, M.G, W.A, R.L, G.S have filed a provisional patent based on this work.

## Author contributions

K.Y.V. and A.K. contributed equally to this work. G.S. and R.L. conceptualized the idea and directed the work. V.K.Y., A.K., N.D.P, G.S. and R.L. designed the experiments. A.K. and W.A. performed synthesis of molecules. K.Y.V., N.D.P., J.W. and K.D.M.D. established animal models. K.Y.V., N.D.P., A.J., D.K.S., M.G., N.T.V.M., A.R. and J.W. performed the experiments. K.Y.V., N.D.P., A.K., A.J., D.K.S., N.T.V.M., G.S, R.L. analyzed the data. K.Y.V., A.K, R.L., and G.S. wrote the article with contribution of all authors. All authors read and approved the final article.

