## Supplementary material for "Synthesis of ionizable lipopolymers using split-Ugi reaction for pulmonary delivery of various size RNAs and gene editing": Supporting information.docx

* Both authors contributed equally

#co-corresponding authors

**Table S1**: Composition of Ugi modified polymers. The percentage value refers to the targeted modification degree. Samples U1-U10 are highlighted as were synthesized under a slightly different synthetic protocol.

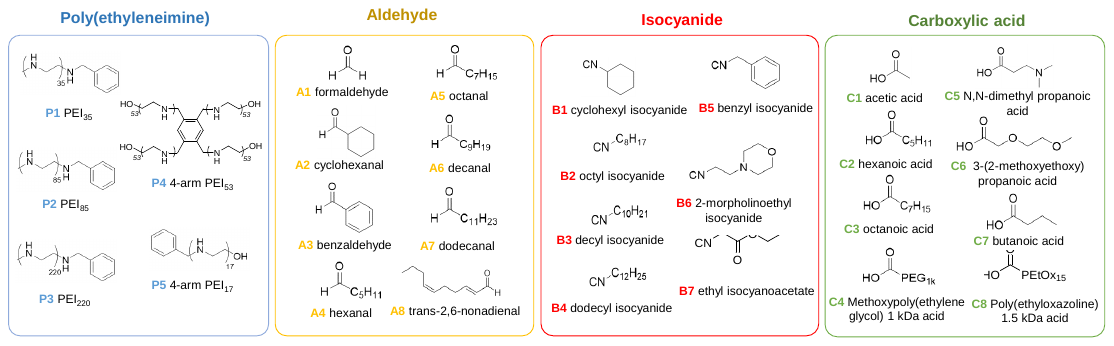

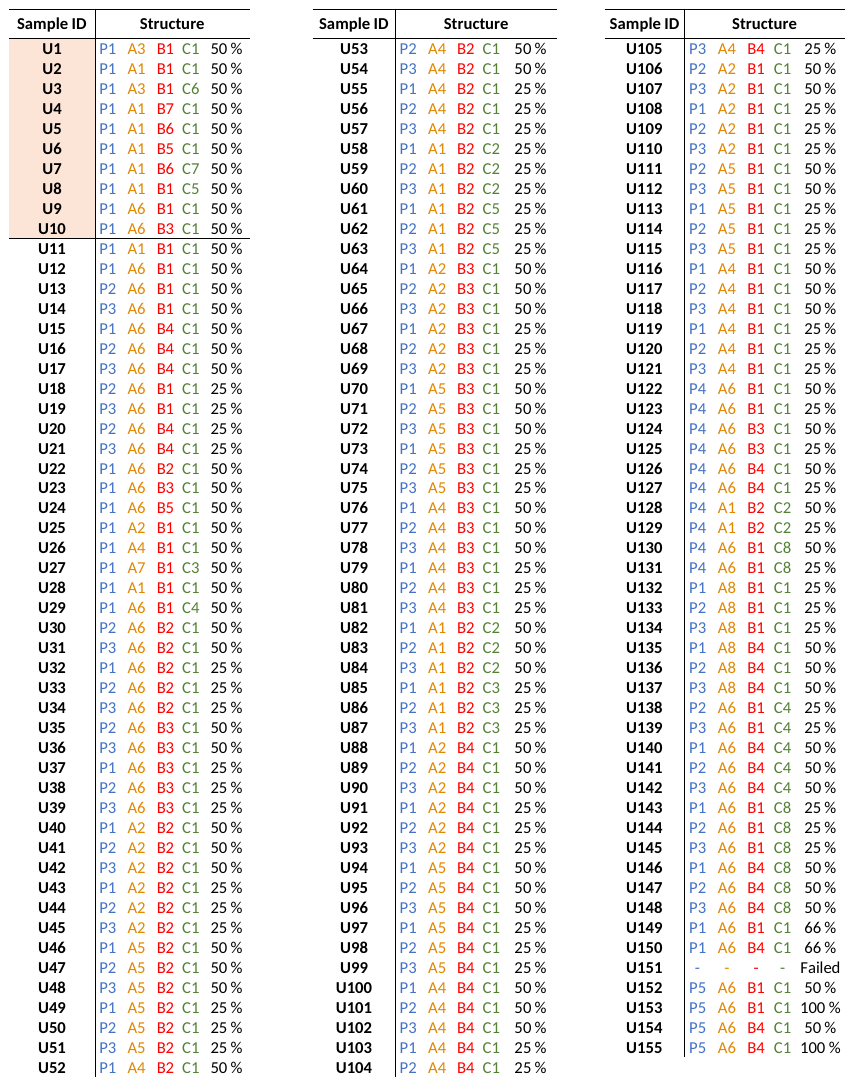

**Fig. S1. a)** ^1^H NMR spectra in MeOD of PEI35 modified with decanal / decylisocyanide / acetic acid (U15, P_1_A_6_B_4_C_1_-50%). Here the signal ‘g’ corresponds to both the CH3 group of the aldehyde and isocyanidederived moeities. The integral of the methyl amide shows a modification of only 14 %, lower than the targeted 25 %. Such a result was quite common across the sample library, and it appears a portion of the acid remains unreacted as can be seen by the signals for residual free acetic acid; **b)** DMF SEC chromatograms of a range of Ugi modified samples derived from the PEI35 backbone; **c)** ^1^H NMR spectra in MeOD of lead compound U155. Modification degree was calculated by comparing integral ‘a’ of phenyl end group to integral ‘i’ from CH3 units of the aldehyde and isocyanide components **d)** CHCl_3_ SEC chromatogram of U155.

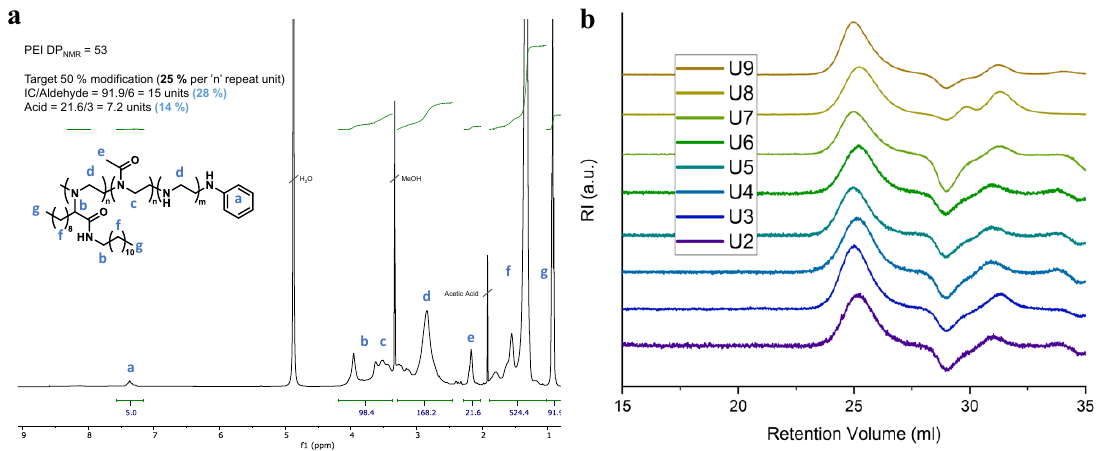

**c**

**d**

**Fig. S3. a-b)** In vivo bioluminescent images of BALB/c mice 5-6 h **(a)** and 24 h **(b)** post I.V. injection of U155 polyplexes encapsulated 5 μg Fluc mRNA per mouse; **c)** ex vivo bioluminescent images of BALB/c mice 24 h (post I.V. injection of U155 polyplexes encapsulated 5 μg Fluc mRNA per mouse; **d)** ex vivo bioluminescent images of BALB/c mice 5-6 h (post I.V. injection of large (app. 600 nm) U155 polyplexes encapsulated 5 μg Fluc mRNA per mouse.

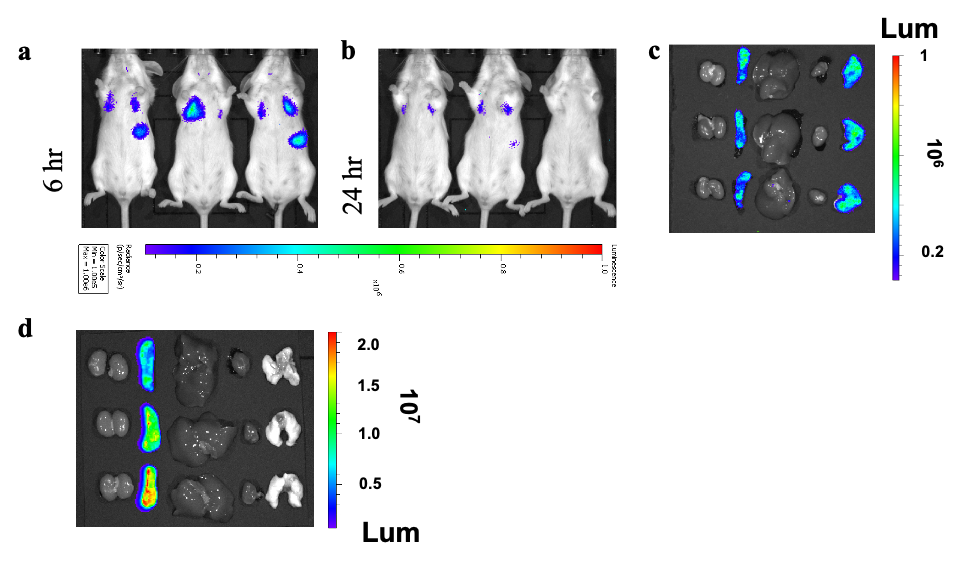

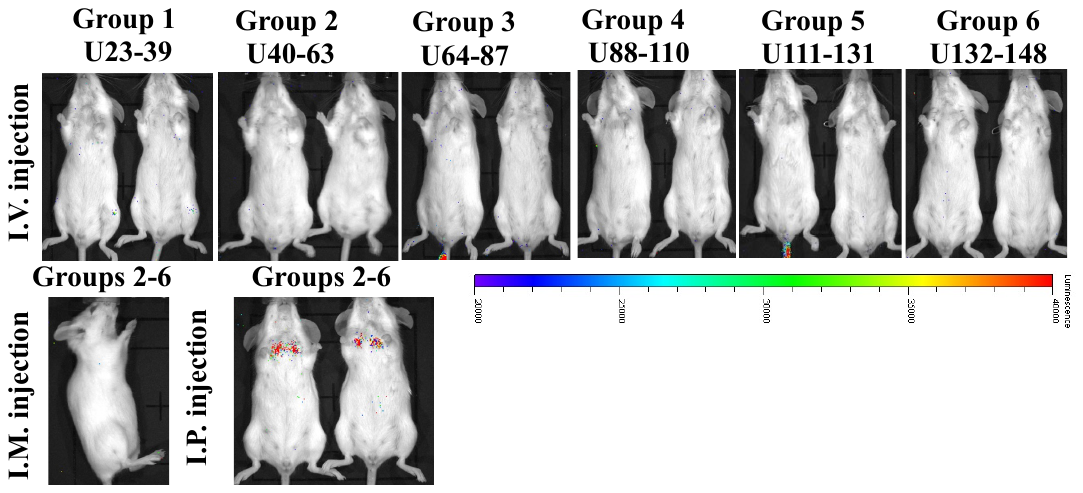

**Fig. S2**. In vivo screening of grouped Fluc mRNA-loaded polyplexes. Representative IVIS images of BALB/c mice 5-6 h following polyplexes administration. For in vivo screening studies, results were obtained from two mice per group. Each particles type in the group was injected at the dose of 1 μg Fluc mRNA/mouse.

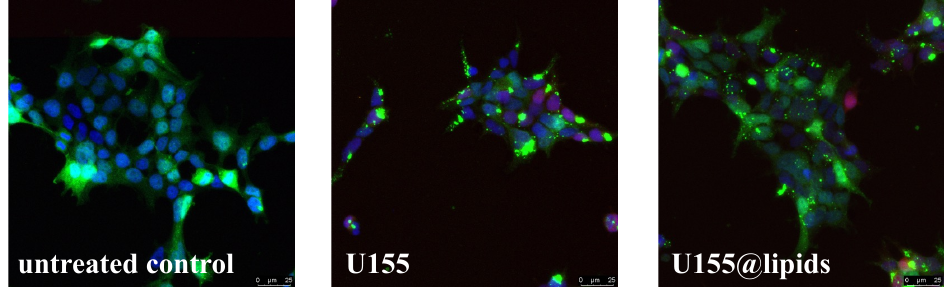

**Fig. S5.** Representative images of Gal9-HEK293 cells after 24 h incubation with mCherry RNA- loaded nanoparticles. mCherry RNA dose 150 ng per well.

**Fig. S4.** DSPG content screening. **a)** primary trachea cells co-cultured with fibroblast were treated with U155@lipids with various soyPC/DSPG ratios and encapsulated Fluc mRNA. The relative luciferase expression (normalized to cell viability fluorescent signal) after 24 h incubation with polyplexes is shown (2 biological replicates with 6 technical replicates). Fluc mRNA dose 200 ng per well. **b)** In vivo bioluminescent images of BALB/c mice 5-6 h post I.V. injection of 2 μg Fluc mRNA per mouse encapsulated in U155@lipids with 22 mol % (2.75x10^6^ total efflux) or 31.5 mol % DSPG (1.01x10^6^ total efflux).

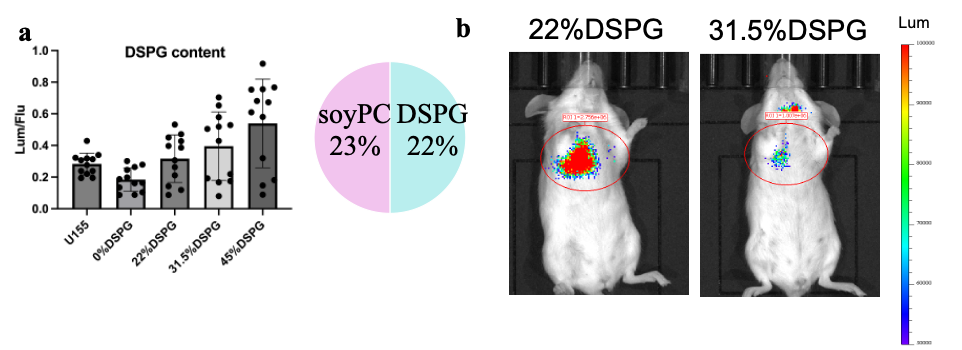

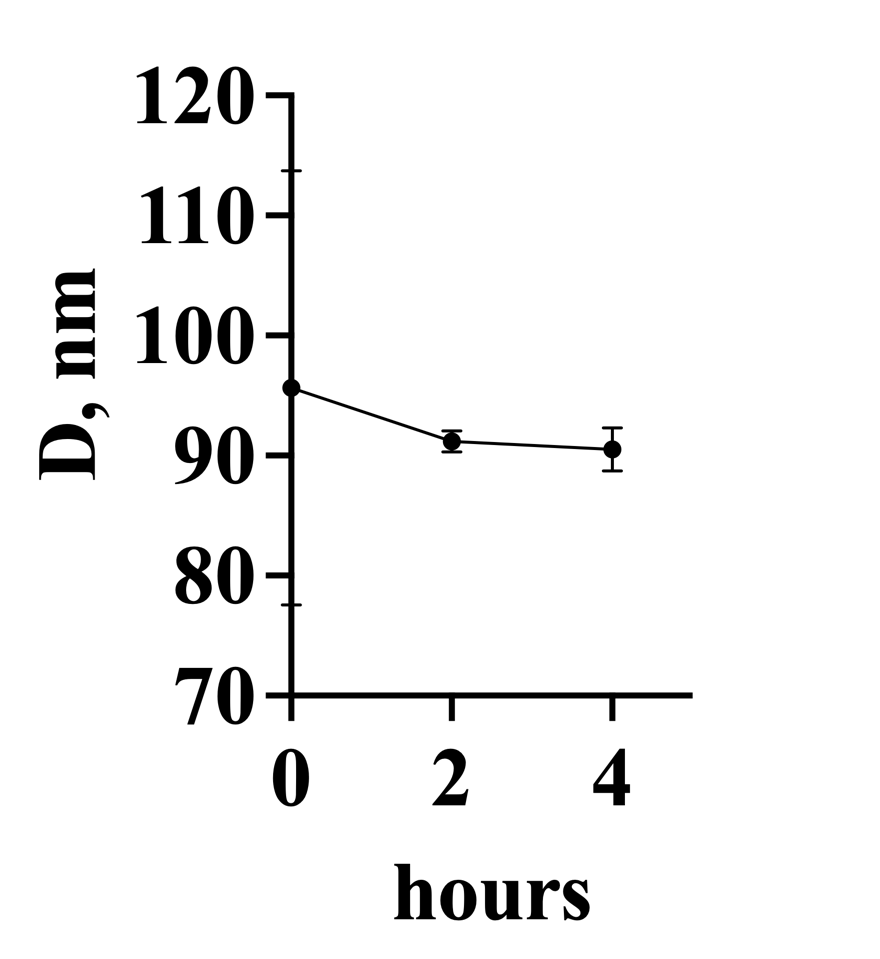

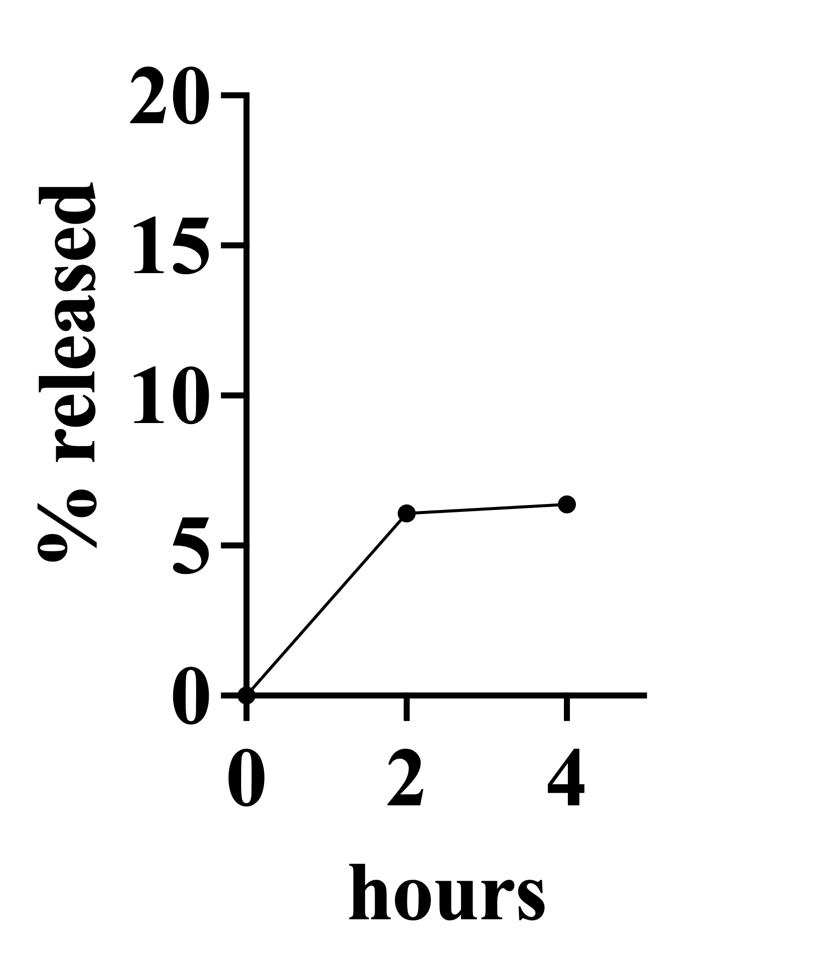

**a**

**b**

**Fig. S7. a)** diameter of DiD-labeled U155@lipids nanoparticles during dialysis against 50% serum at 37 ^o^C; **b)** DiD release from U155@lipids nanoparticles during dialysis against 50% serum at 37 ^o^C. Data are presented as Mean ± SD (n=3).

**Fig. S6.** ex vivo bioluminescent images of BALB/c mice 5-6 h post I.V. injection of In vivo JetPEI encapsulated 10 μg Fluc mRNA per mouse.

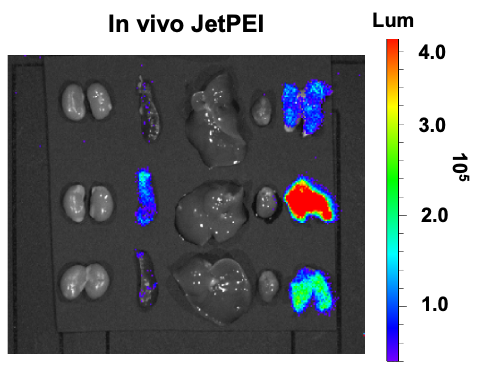

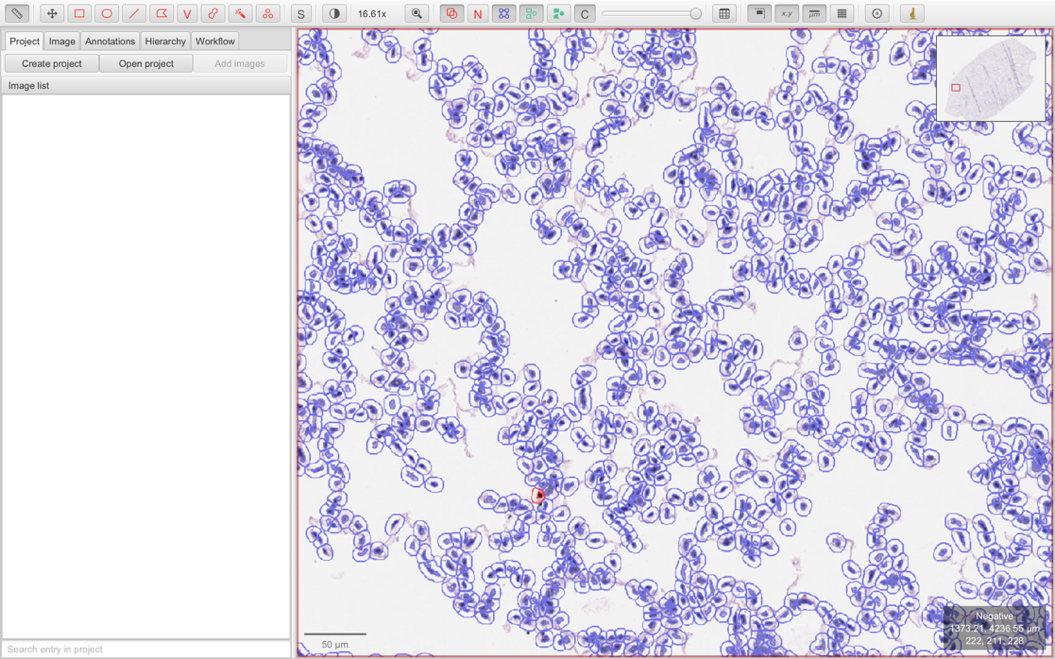

**Fig. S8. a-b)** In vivo bioluminescent images of BALB/c mice 5-6 h (a) and 24 h (b) post I.V. injection of U155@lipids encapsulated 5 μg Fluc mRNA per mouse; **c)** ex vivo bioluminescent images of BALB/c mice 24 h post I.V. injection of U155@lipids encapsulated 5 μg Fluc mRNA per mouse.

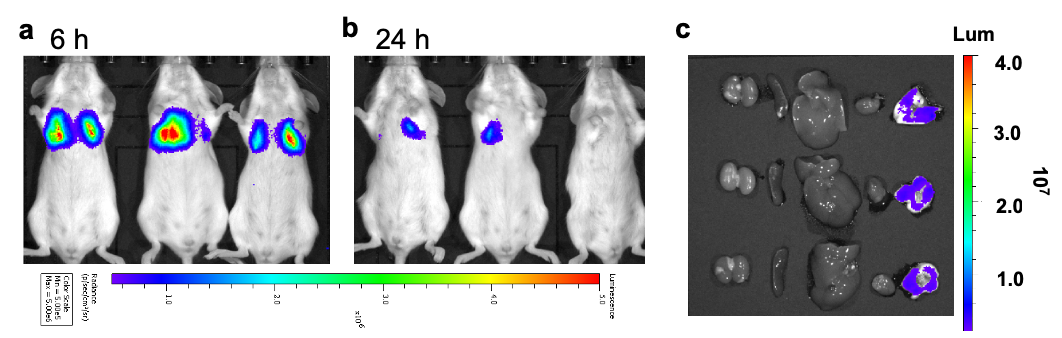

**Fig. S9. a-b)** In vivo bioluminescent images of BALB/c mice 5-6 h (a) post I.V. injection of U155@lipids encapsulated total 5 μg Fluc mRNA per mouse. a) 1^st^ dose of I.V. injected U155@lipids encapsulated 5 μg Fluc mRNA; **b)** 2^nd^ dose of I.V. injected U155@lipids 5 μg Fluc mRNA. Mice were treated 48 h post 1^st^ dose injection.

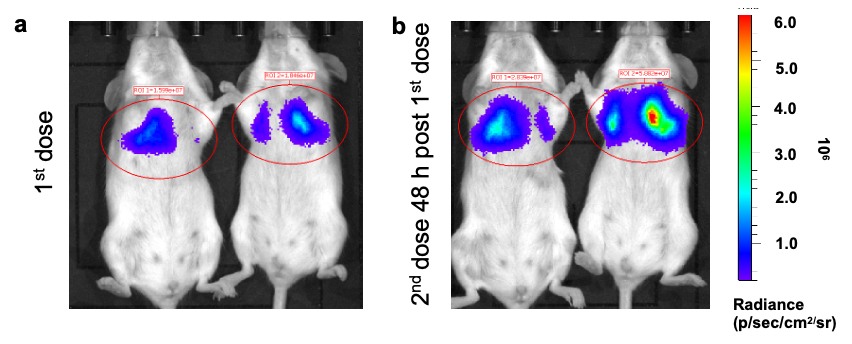

**Fig. S10.** Representative image of how QuPath’s tool select nuclei.

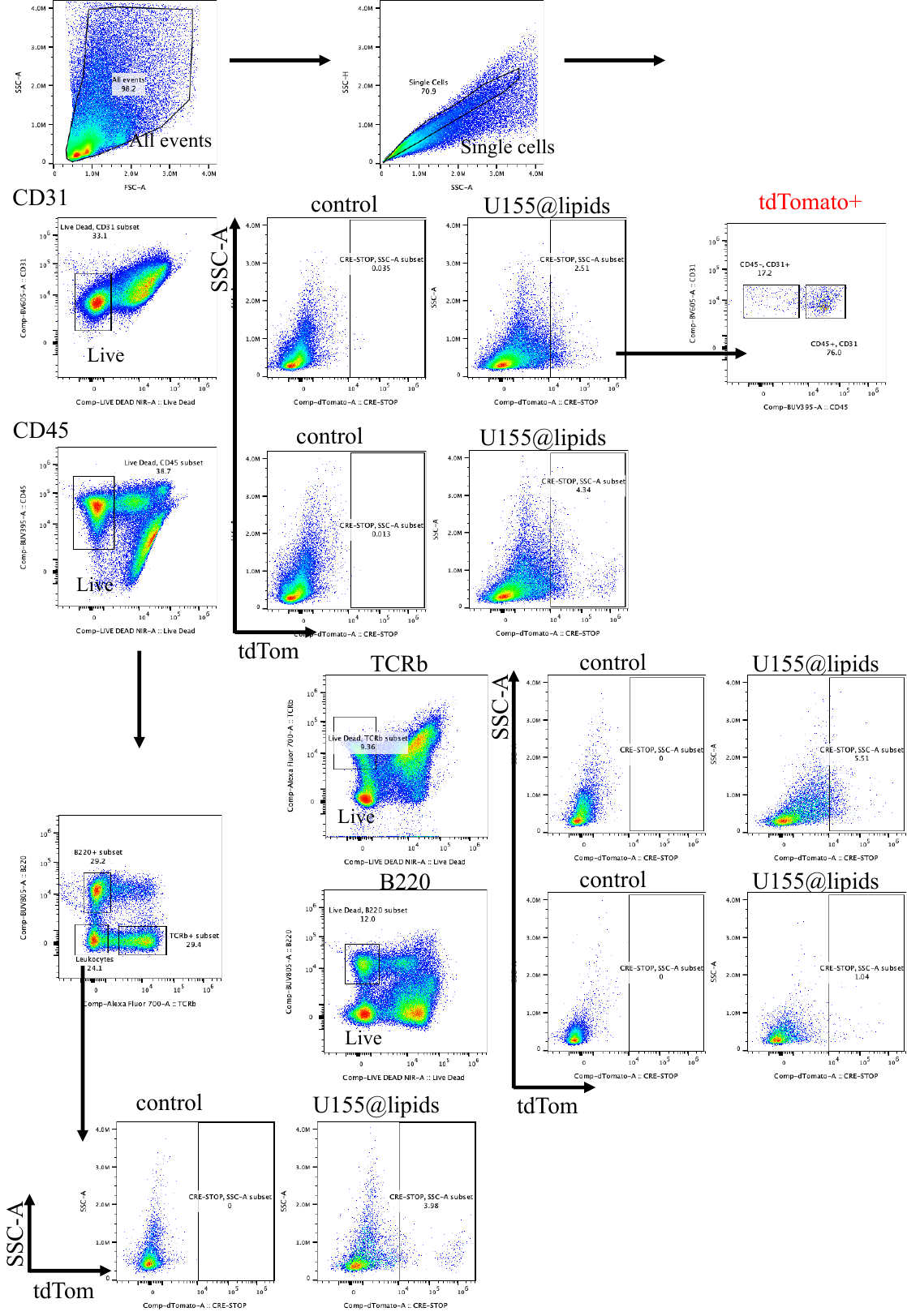

**Fig. S11.** Workflow of flow cytometry gating strategy for lungs tissue.

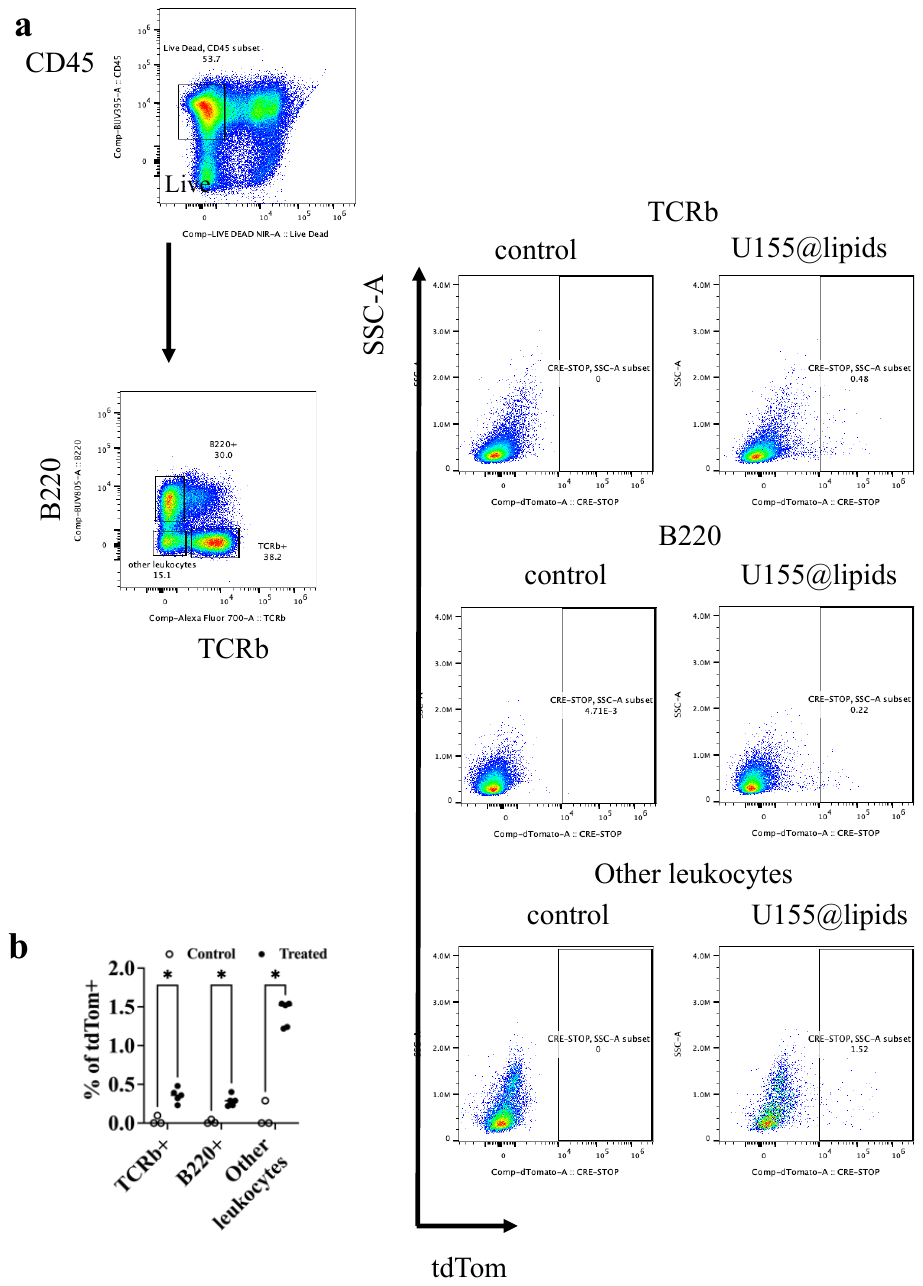

**Fig. S12. a)** Workflow of flow cytometry gating strategy for spleen tissue; **b)** transfection of immune cells of spleen after I.V. injection of 10 ug Cre mRNA encapsulated in U155@lipids in Ai9 mice. Data are presented as Mean ± SD (n=3-5).

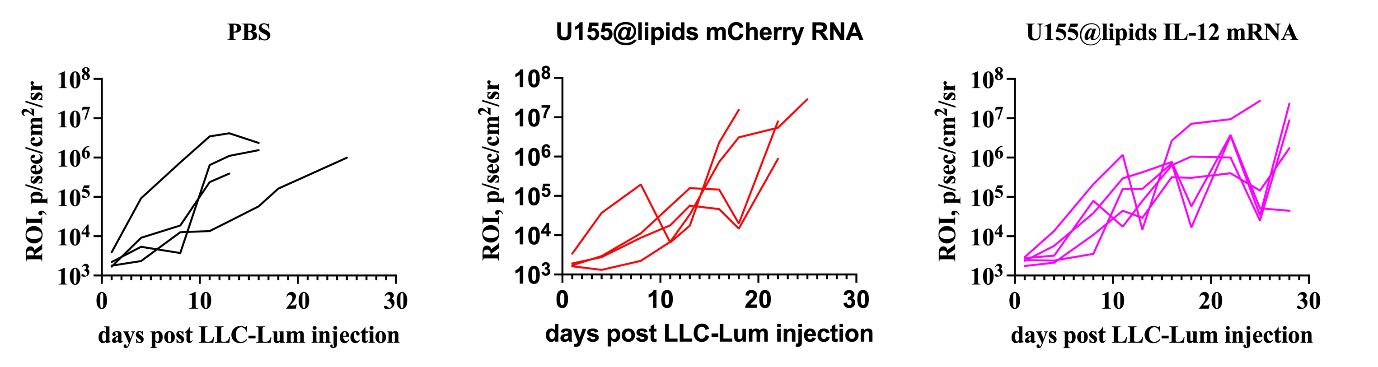

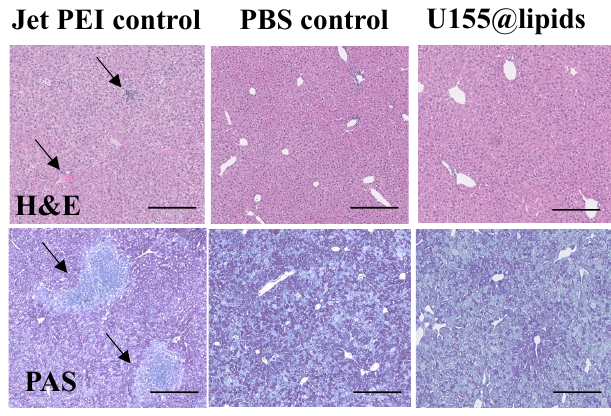

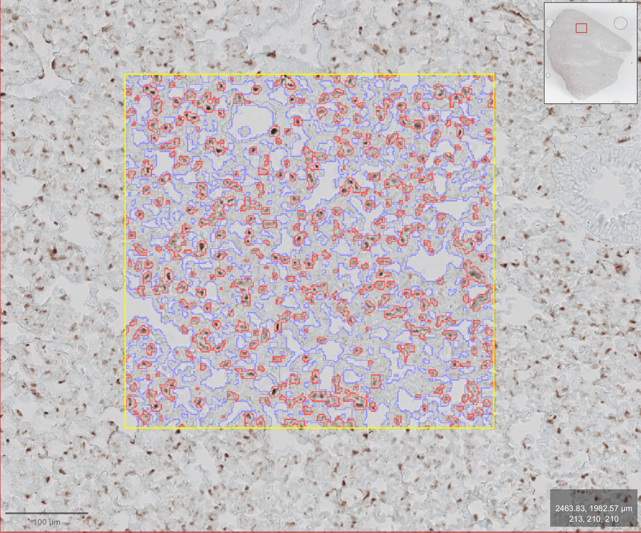

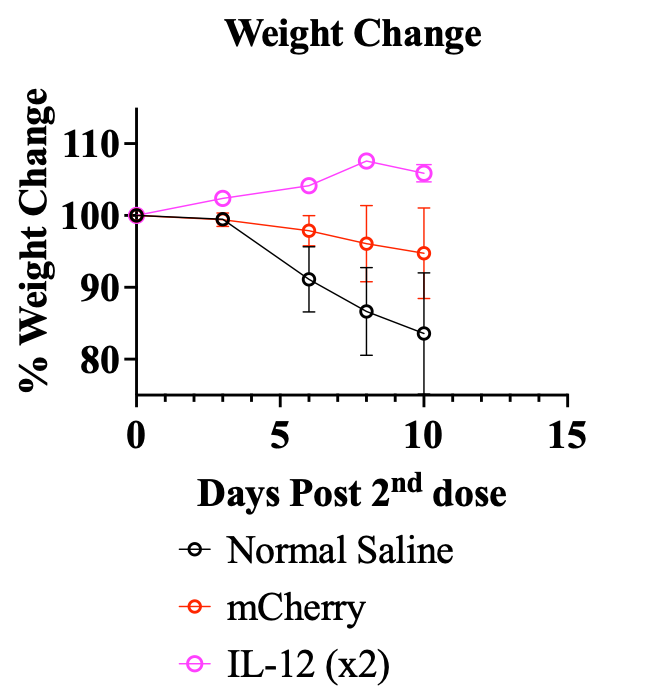

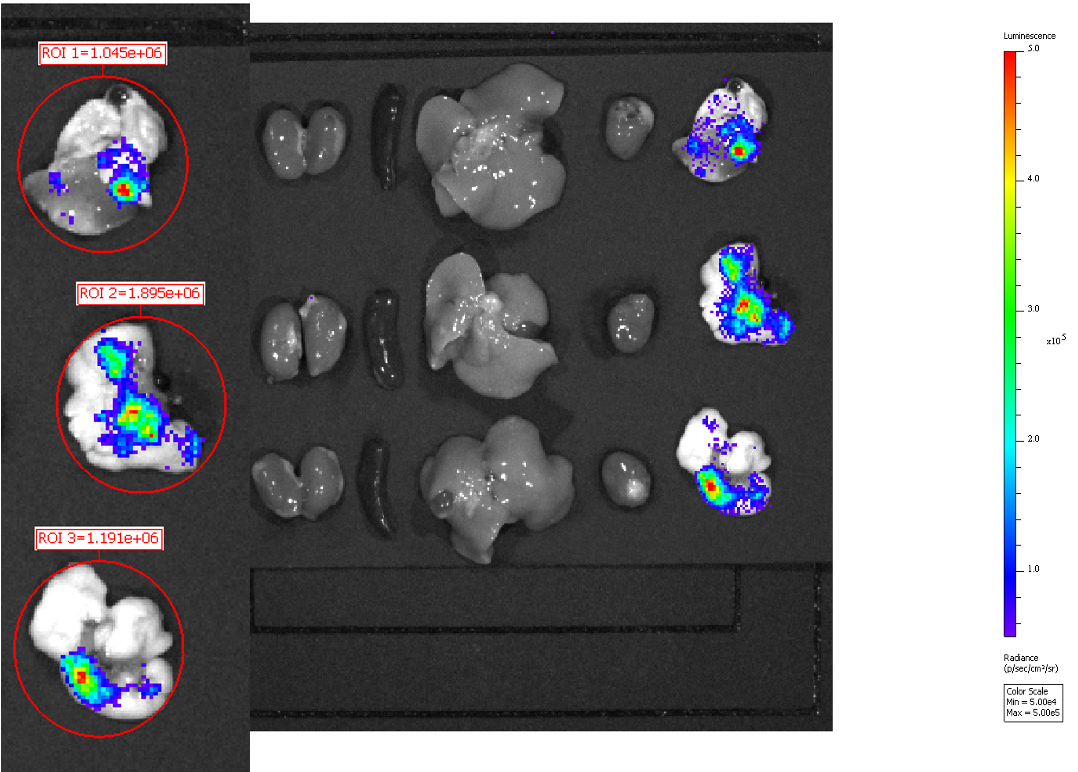

**Fig. S15.** Individual tumor growth over time for each treatment group. n=4 biologically independent mice for PBS and mCherry RNA groups, n=5 biologically independent mice for IL-12 mRNA group.

**Fig. S14.** Representative tissue images of liver stained with H&E and PAS-fast green. Black arrows show tissue damage. Scale bar is 200 μm.

**Fig. S13.** Representative image of how QuPath’s tool select tdTomato+ stained areas (red) and tissue (blue).

**Fig. S16.** Body weight change, normalized to weight 24 hr post last treatment, over time for mice in each treatment group. n=4 biologically independent mice for PBS and mCherry RNA groups, n=5 biologically independent mice for IL-12 mRNA group.

**Fig. S17.** *ex vivo* Fluc mRNA transfection by polymer-lipid U155@lipids nanoparticles. Bioluminescent images of BALB/c mice after treatment of 2 μg Fluc mRNA per mouse delivered via 5 hour post-intratracheal installation.

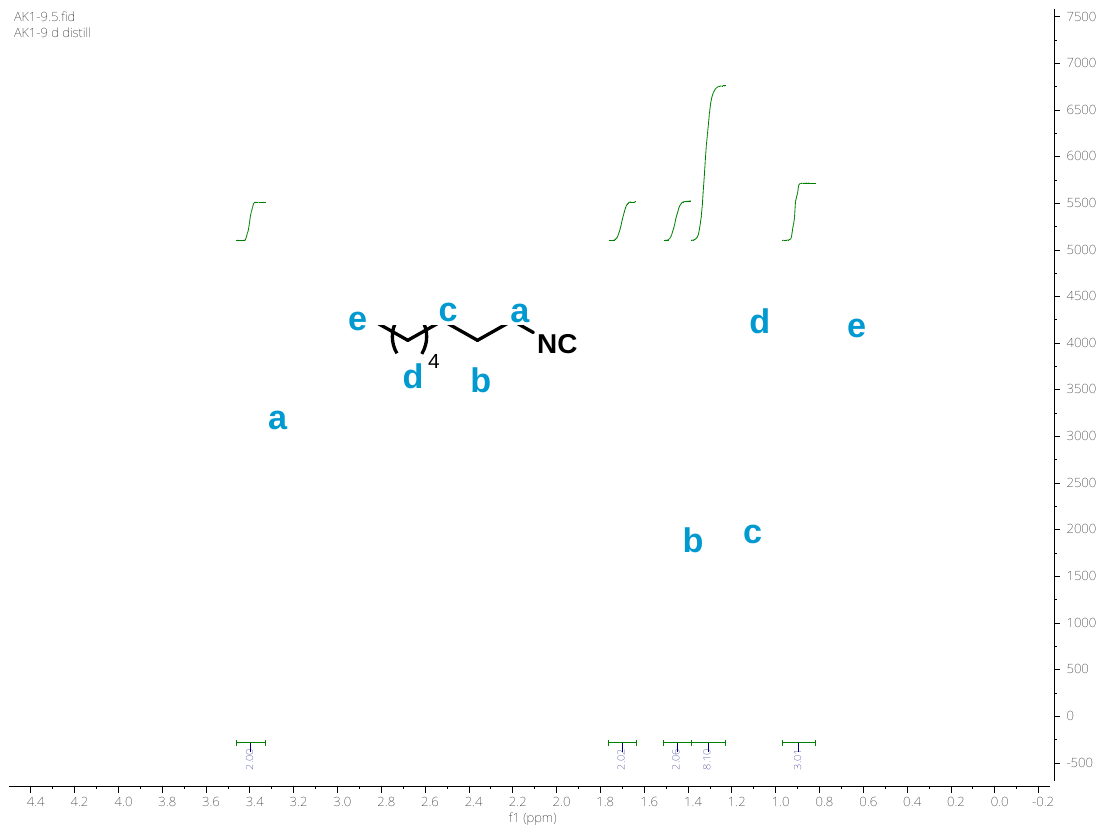

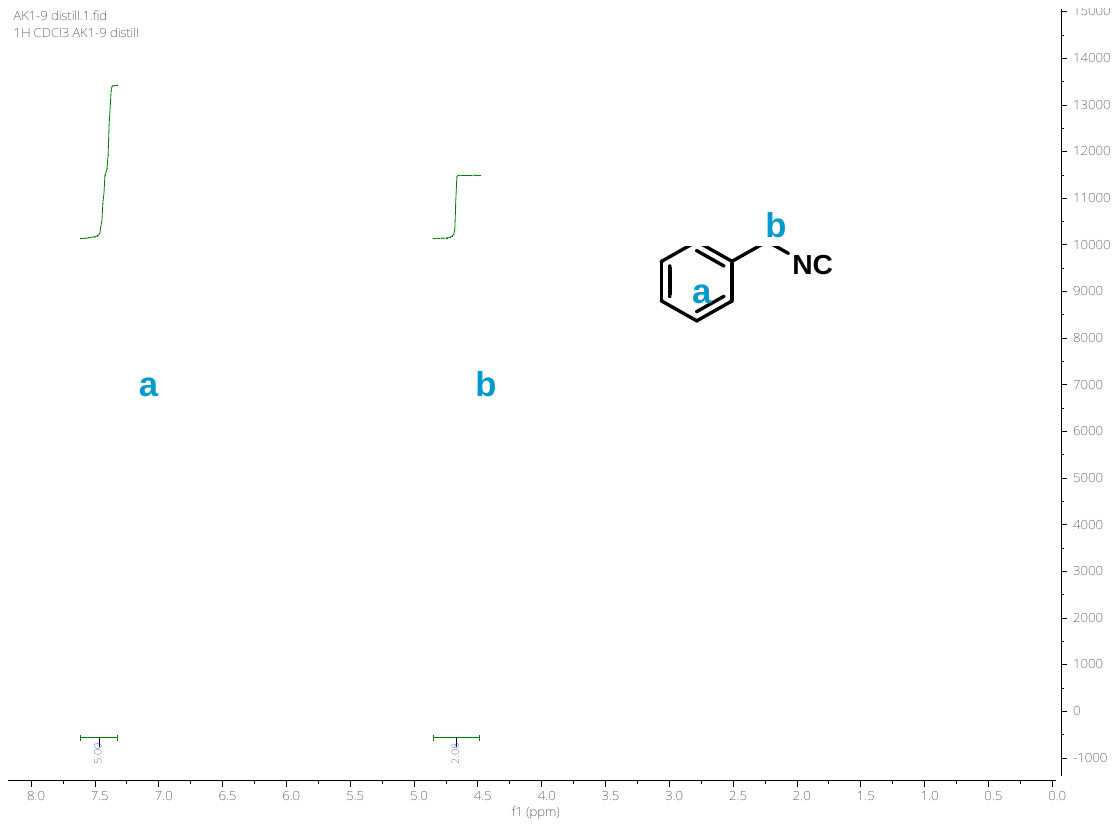

**Fig. S19.** ^1^H NMR spectra in CDCl_3_ of octylisocyanide.

**Fig. S18.** ^1^H NMR spectra in CDCl_3_ of benzoisocyanide.

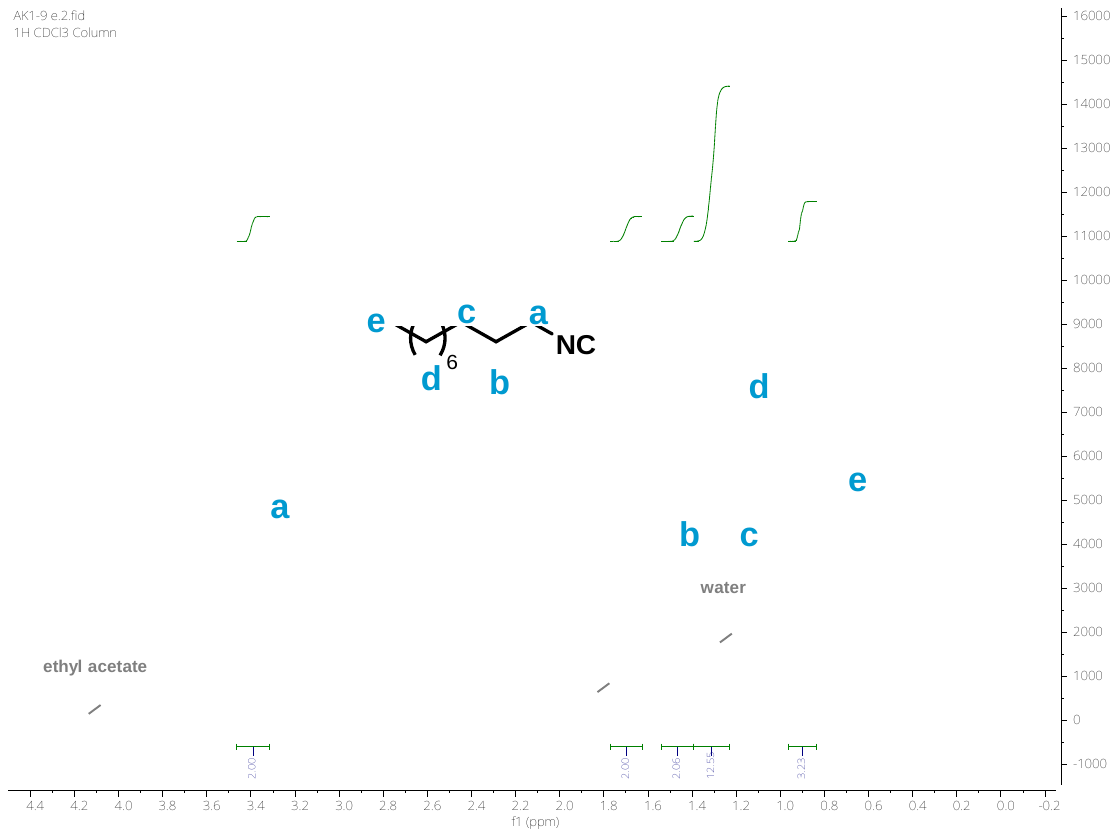

**Fig. S20.** ^1^H NMR spectra in CDCl_3_ of decylisocyanide.

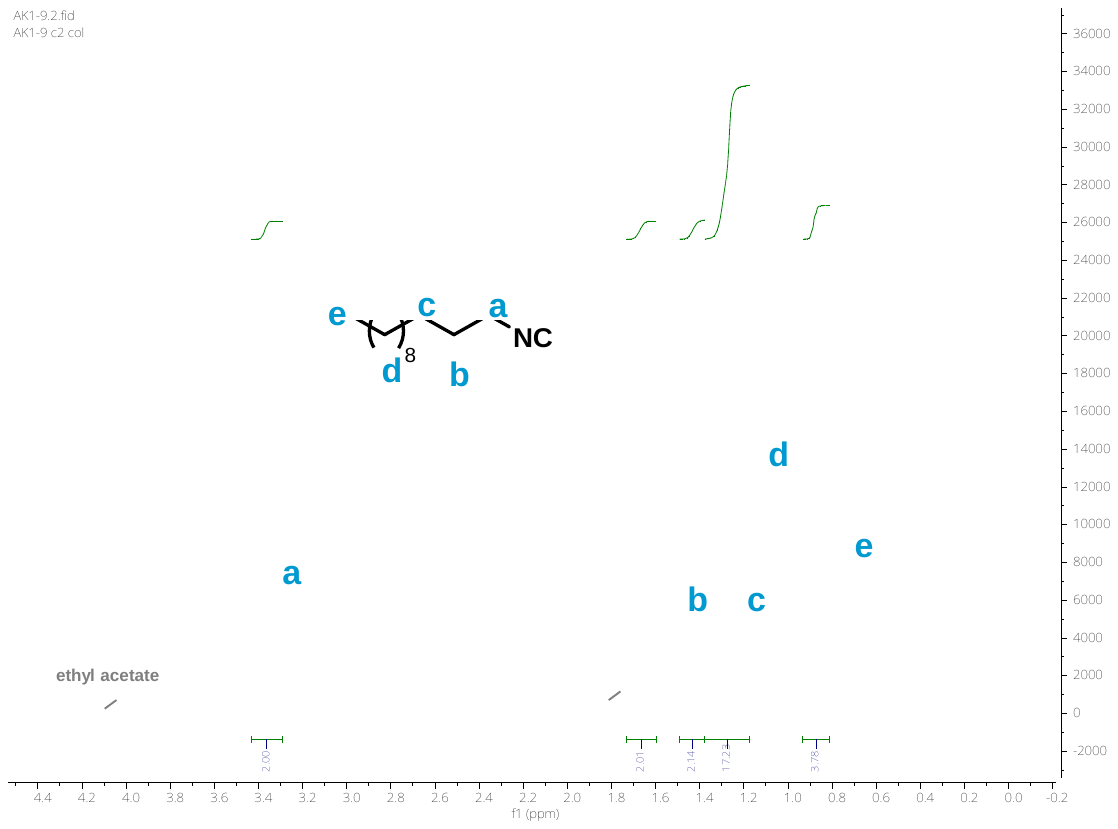

**Fig. S21.** ^1^H NMR spectra in CDCl_3_ of dodecylisocyanide.

**Fig. S22. A** – SEC chromatograms of PEtOx samples of varying molecular weights. **B** – MALDI-ToF spectra of the PEtOx15 sample showing a narrow dispersity and expected chain end functionality.

**Table S2**: Analytical data of the PEtOx polymers. a – degree of polymerization calculated from NMR integration of the phenyl protons (7.3 ppm) compared to the polymer side chain ethyl units (ppm). b – SEC performed in DMF + 0.1% LiBr eluent.

| **Sample** | **Initiator** | **DP_NMR_^a^** | **M_n theo_**  **(g mol^-1^)** | **M_n SEC_**  **(g mol^-1^)^b^** | ***Đ*** |
| --- | --- | --- | --- | --- | --- |
| PEtOx_15_ | BzBr | 17 | 1595 | 1810 | 1.38 |
| PEtOx_35_ | MeOTf | 53 | 3591 | 6470 | 1.13 |
| PEtOx_85_ | MeOTf | 120 | 8547 | 12300 | 1.35 |
| PEtOx_220_ | MeOTf | - | 21930 | 25000 | 1.54 |

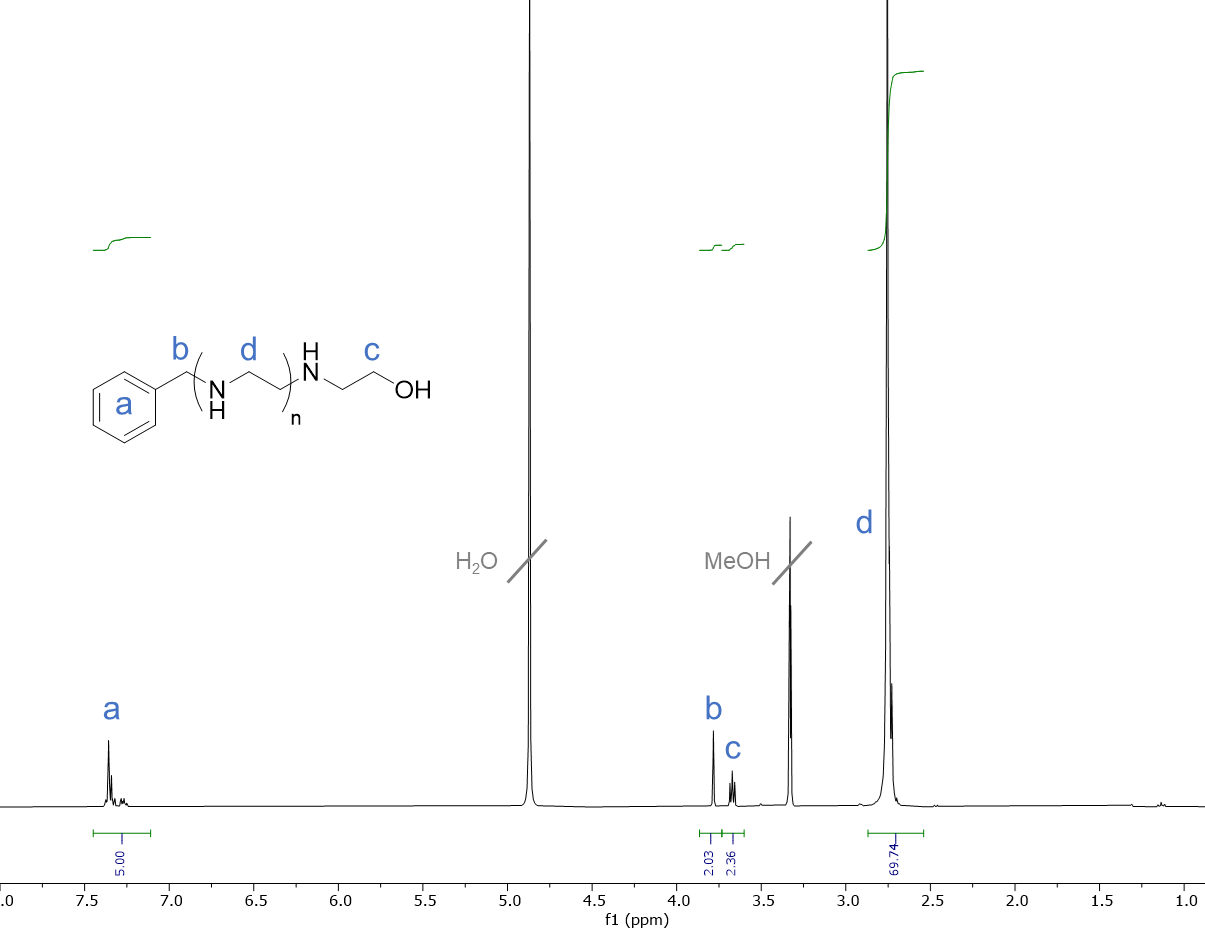

**Fig. S23.** ^1^H NMR of poly(ethyleneimine) in MeOD prepared from hydrolysis of the PEtOx_15_ sample, NMR integrals consistent with a degree of polymerization = 17.

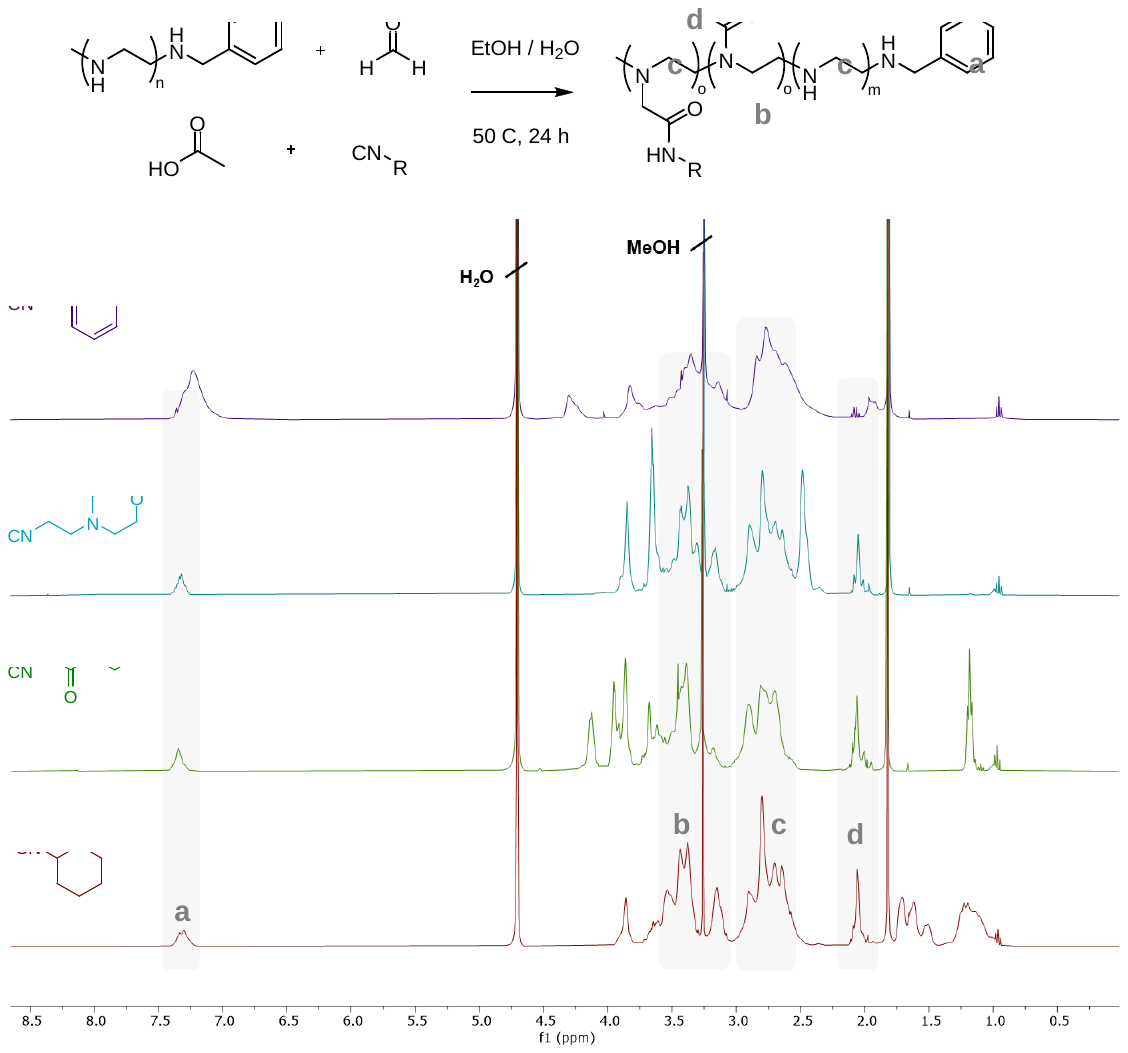

**Fig. S24.** ^1^H NMR spectra in MeOD of PEI Ugi derivatives, functionalized with formaldehyde/acetic acid and varying isocyanide reagents. The core polymer backbone peaks are similar across all four samples, but additional peaks from the isocyanide moiety are identifiable in each case.

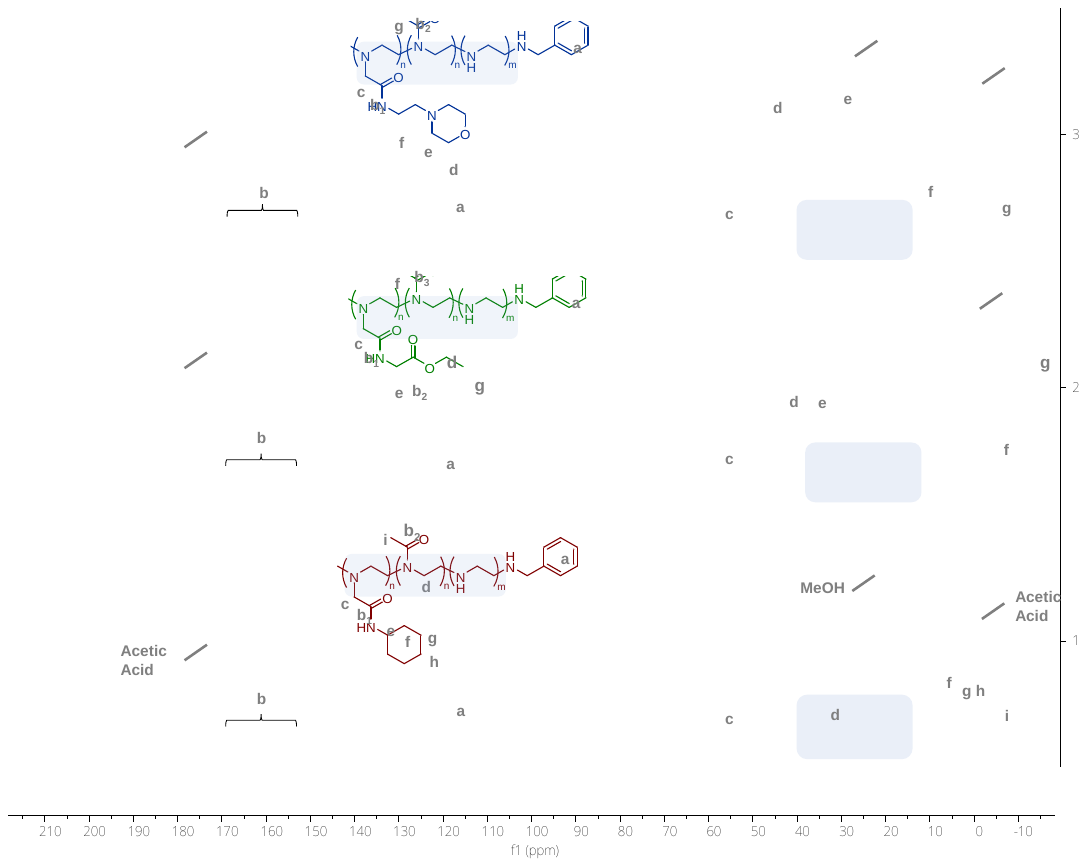

**Fig. S25.** ^13^C NMR spectra in MeOD of PEI ugi derivatives. Assignments made in combination with HSQC analysis.

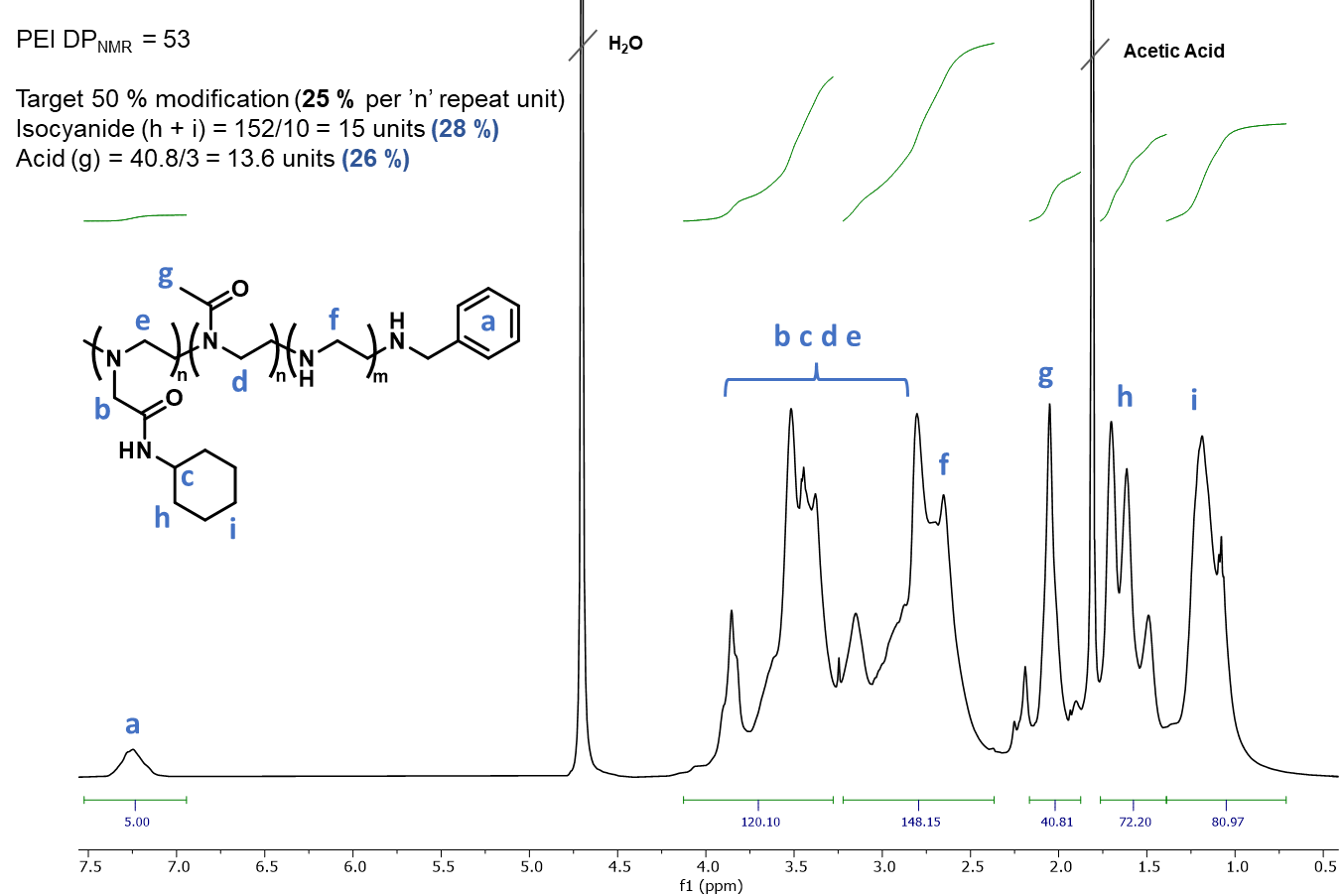

**Fig. S26.** ^1^H NMR spectra in MeOD of a PEI_35_ modified with formaldehyde / cyclohexylisocyanide / acetic acid. Modification % were calculated assuming the polymer degree of polymerization is the same as the PEI starting material (DP 53). The integral of the phenyl end group protons (a) is then compared to the alkyl protons (ppm 0.9 – 1.7) of the cyclohexyl unit to calculate the number of isocyanide moieties per polymer chain. The methyl amide derived from reaction with acetic acid is calculated from the signal at 2 ppm (g). Targeting a total modification of 50 % through the split Ugi reaction should result in a final composition of 50 % unreacted PEI repeat unit and 25 % each of the tertiary amine and amide repeat units. In this example values of 28 / 26 % are obtained, consistent with the theoretical considering error in NMR integration. Signal ‘b’ overlaps with polymer backbone signals so the number of aldehyde moieties introduced cannot be calculated, but according to the Ugi mechanism an equimolar quantity to the isocyanide reagent can be expected.

:

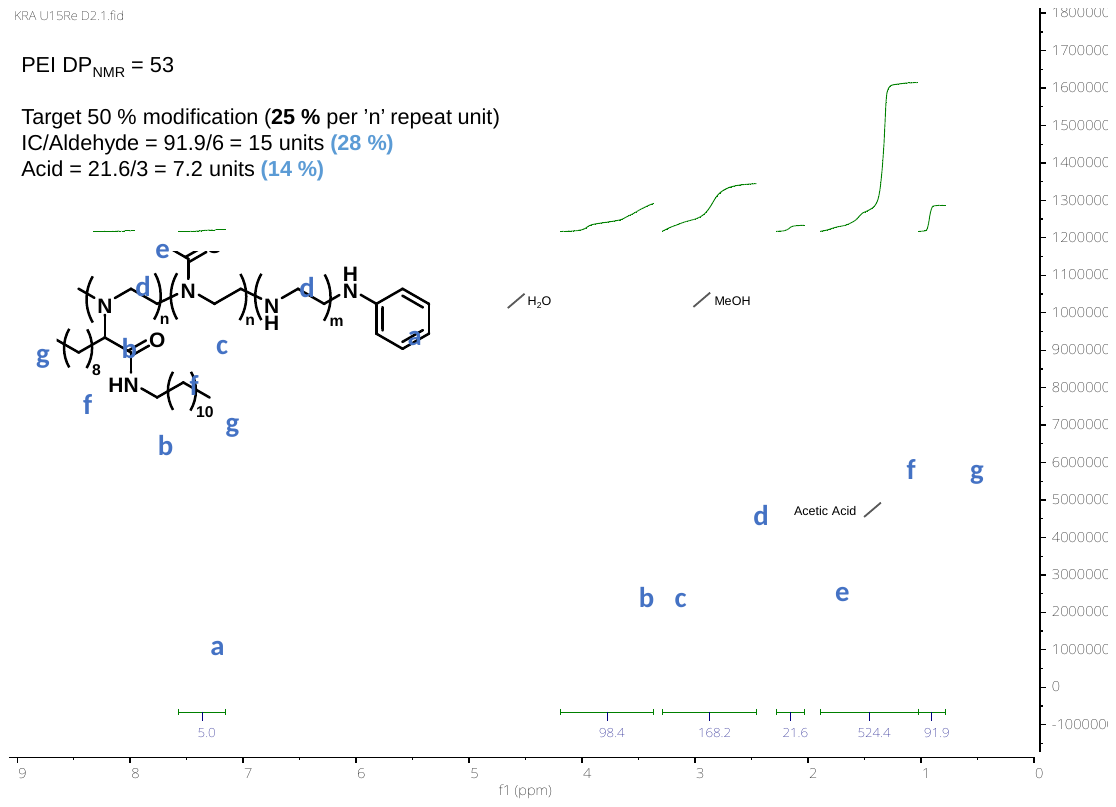

**Fig. S27**: ^1^H NMR spectra in MeOD of PEI_35_ modified with decanal / decylisocyanide / acetic acid. Modification % was calculated using the same method as described above (Figure S). Here the signal ‘g’ corresponds to both the CH_3_ group of the aldehyde and isocyanide derived moeities. The integral of the methyl amide shows a modification of only 14 %, lower than the targeted 25 %. Such a result was quite common across the sample library, and it appears a portion of the acid remains unreacted as can be seen by the signals for residual free acetic acid.

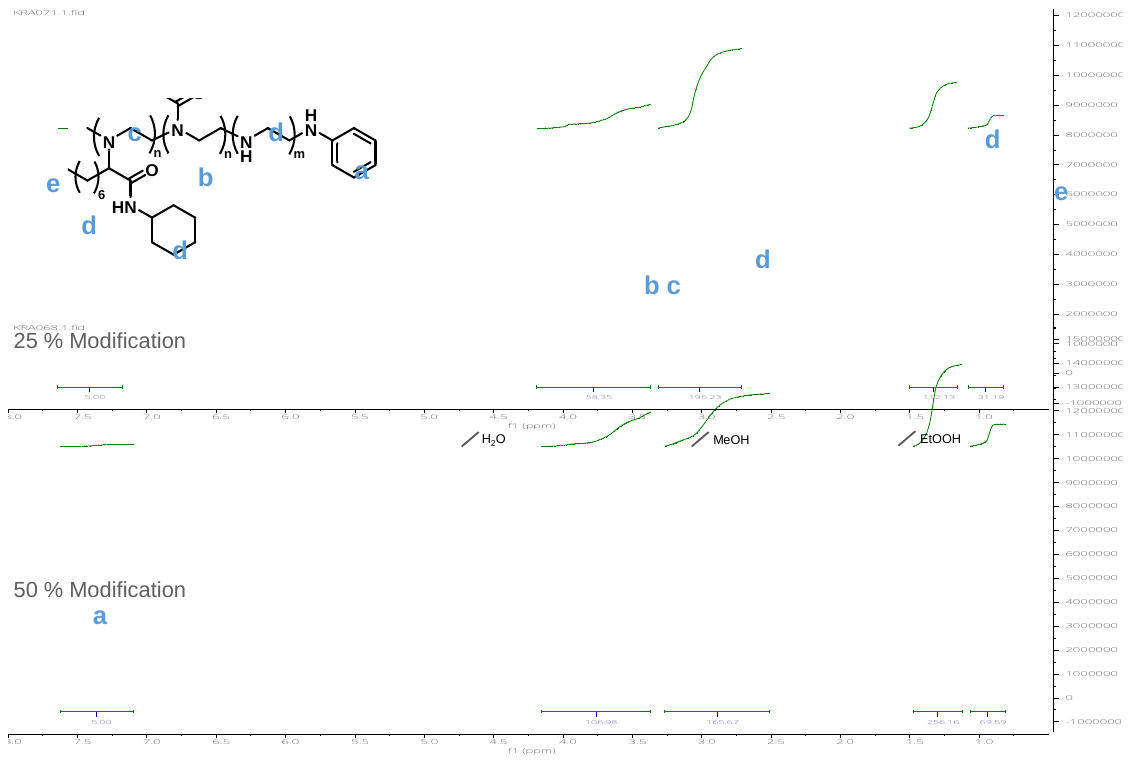

**Fig. S28**: ^1^H NMR spectra in MeOD of a PEI_35_ modified with octanal / cyclohexylisocyanide / acetic acid at two different modification densities. Integration shows signals from the alkyl chains of aldehyde/isocyanide derived species (d,e) are approximately doubled for 50% modification over 25 %, as expected. The total integration of the polymer backbone regions (b, c, d) are similar in both samples, but the PEI repeat unit at 2.8 ppm is reduced for 50 % modification sample indicating a larger portion has been successfully reacted.

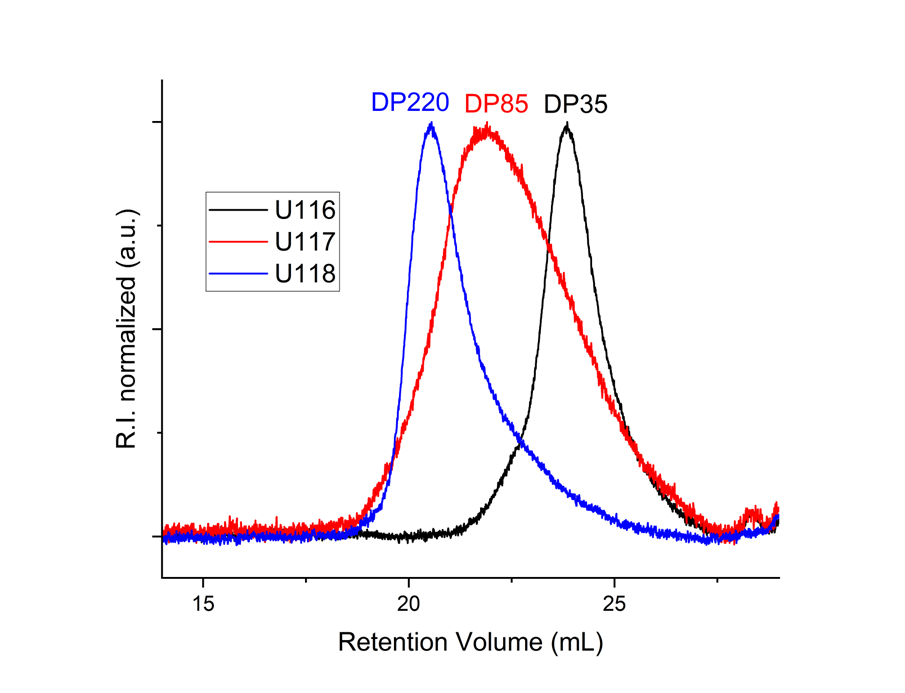
**Fig. S29**: CHCl_3_ SEC chromatograms of samples U116 (P_1_A_4_B_1_C_1_-50%), U117 (P_2_A_4_B_1_C_1_-50%), U118 (P_3_A_4_B_1_C_1_-50%). These samples were modified with the same reagents but derived form different length of PEI backbones, the DP indicated above the corresponding traces.

**Effect of equivalents of carboxylixc acid reagent**

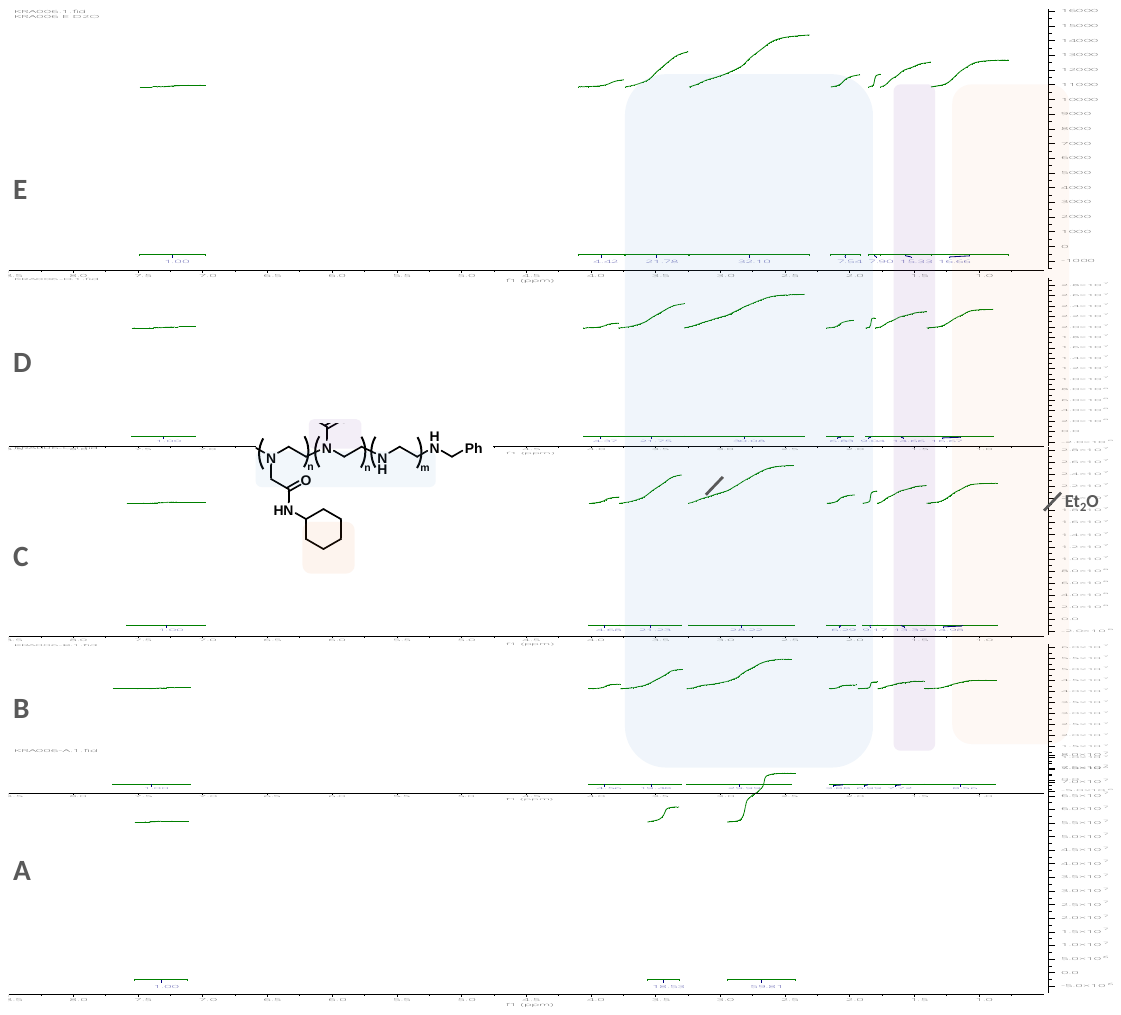
**Fig. S30**: ^1^H NMR spectra in MeOD of PEI-Ugi derivatives synthesized with varying quantities of the acid component, see table S for equivalents of acid in each experiment. Polymers were precipitated three times into diethyl ether and dried under vacuum before analysis. A trend of higher functionalization of cyclohexyl units with higher equivalents of acid used in the reaction was observed. For entry A, where no acid reagent is used, the Ugi reaction fails and no isocyanide units are successfully reacted, likely only imine formation with the aldehyde occurs under these conditions.

**Table S3**: Reaction conditions and obtained modification as determined by NMR for Ugi reactions with varying equivalents of the carboxylic acid component, NMR spectra shown above in Figure S. The equivalents of acid were calculated taking the PEI repeat unit as 1 eq. A total modification of 50 % was targeted, meaning an expected functionalization of 25 % for the acid and aldehyde/isocyanide units respectively. A ratio of reagents 1 : 0.25 : 0.25 : X (PEI : aldehyde : isocyanide : acid) was used, were X is shown in the table. Entry B uses the minimum stoichiometry of acid needed for the Ugi, however, this leads to sub-quantitative functionalization. This presumably arises from ionic interactions of the acid with excess secondary amine units of the PEI backbone, reducing the reaction rate. Increasing the acid to a slight excess (entry C) leads to a significant increase in modification, and further modest increase is seen using higher amounts (entries D/E). As there was no apparent downside of using the highest acid concentration, this was selected as the standard condition for further library synthesis.

| **Exp.** | **Eq. Acid** | **Acid modified units** | **Isocyanide modified units** |
| --- | --- | --- | --- |
| A | 0 | 0.0 % | 0.0 % |
| B | 0.25 | 12.2 % | 14.6 % |
| C | 0.375 | 19.8 % | 25.1 % |
| D | 0.5 | 21.5 % | 27.7 % |
| E | 1 | 23.7 % | 28.9 % |

**Table S4:** Guide RNA, primers, and cycling conditions used for editing studies.

| **S.N.** | **Primer Name** | **Sequence (5' -> 3')** |
| --- | --- | --- |
| 1. | Ai9_NGS_F1_F | ACACTCTTTCCCTACACGACGCTCTTCCGATCTATACGAAGTTATTCGCGATG |
| 2. | Ai9_NGS_F2_F | ACACTCTTTCCCTACACGACGCTCTTCCGATCTGTTGTGGTTTGTCCAAAC |
| 3. | Ai9_NGS_R | GACTGGAGTTCAGACGTGTGCTCTTCCGATCTTGTTTCAGGTTCAGGGGGAG |
| 4. | PD1_NGS _F | ACACTCTTTCCCTACACGACGCTCTTCCGATCTttgccttggggtgcagggag |
| 5. | PD1_NGS _R | GACTGGAGTTCAGACGTGTGCTCTTCCGATCTgtgtcagagggagcaaatgc |

**Table S5:** Thermocycling methods for PCR for NGS library preparation.

|  | **1^st^ PCR Ai9** | | **1^st^ PCR PD1** | | **2^nd^ PCR** | |
| --- | --- | --- | --- | --- | --- | --- |
| **Step** | **Temp.** | **Time** | **Temp.** | **Time** | **Temp.** | **Time** |
| Initial Denaturation | 95.0 °C | 3 min | 95.0 °C | 3 min | 98.0 °C | 2 min |
| 30 Cycles:  Denaturation  Annealing  Extension | 98.0 °C  60.0 °C  72.0 °C | 20 sec  20 sec  30 sec | 98.0 °C  65.0 °C  72.0 °C | 20 sec  20 sec  30 sec | 98.0 °C  62.0 °C  72.0 °C | 10 sec  20 sec  30 sec |
| Final Extension | 72.0 °C | 30 sec | 72.0 °C | 30 sec | 72.0 °C | 2 min |

IL-12 mRNA sequence

ATGTGTCCTCAGAAGCTAACCATCTCCTGGTTTGCCATCGTTTTGCTGGTGTCTCCACTCATGGCCATGTGGGAGCTGGAGAAAGACGTTTATGTTGTAGAGGTGGACTGGACTCCCGATGCCCCTGGAGAAACAGTGAACCTCACCTGTGACACGCCTGAAGAAGATGACATCACCTGGACCTCAGACCAGAGACATGGAGTCATAGGCTCTGGAAAGACCCTGACCATCACTGTCAAAGAGTTTCTAGATGCTGGCCAGTACACCTGCCACAAAGGAGGCGAGACTCTGAGCCACTCACATCTGCTGCTCCACAAGAAGGAAAATGGAATTTGGTCCACTGAAATTTTAAAAAATTTCAAAAACAAGACTTTCCTGAAGTGTGAAGCACCAAATTACTCCGGACGGTTCACGTGCTCATGGCTGGTGCAAAGAAACATGGACTTGAAGTTCAACATCAAGAGCAGTAGCAGTTCCCCTGACTCTCGGGCAGTGACATGTGGAATGGCGTCTCTGTCTGCAGAGAAGGTCACACTGGACCAAAGGGACTATGAGAAGTATTCAGTGTCCTGCCAGGAGGATGTCACCTGCCCAACTGCCGAGGAGACCCTGCCCATTGAACTGGCGTTGGAAGCACGGCAGCAGAATAAATATGAGAACTACAGCACCAGCTTCTTCATCAGGGACATCATCAAACCAGACCCGCCCAAGAACTTGCAGATGAAGCCTTTGAAGAACTCACAGGTGGAGGTCAGCTGGGAGTACCCTGACTCCTGGAGCACTCCCCATTCCTACTTCTCCCTCAAGTTCTTTGTTCGAATCCAGCGCAAGAAAGAAAAGATGAAGGAGACAGAGGAGGGGTGTAACCAGAAAGGTGCGTTCCTCGTAGAGAAGACATCTACCGAAGTCCAATGCAAAGGCGGGAATGTCTGCGTGCAAGCTCAGGATCGCTATTACAATTCCTCATGCAGCAAGTGGGCATGTGTTCCCTGCAGAGTCCGATCGGTTCCTGGAGTAGGGGTACCTGGAGTGGGCAGGGTCATACCGGTCTCTGGACCTGCCAGGTGTCTTAGCCAGTCCCGAAACCTGCTGAAGACCACAGATGACATGGTGAAGACGGCCAGAGAAAAGCTGAAACATTATTCCTGCACTGCTGAAGACATCGATCATGAAGACATCACACGGGACCAAACCAGCACATTGAAGACCTGTTTACCACTGGAACTACACAAGAACGAGAGTTGCCTGGCTACTAGAGAGACTTCTTCCACAACAAGAGGGAGCTGCCTGCCCCCACAGAAGACGTCTTTGATGATGACCCTGTGCCTTGGTAGCATCTATGAGGACTTGAAGATGTACCAGACAGAGTTCCAGGCCATCAACGCAGCACTTCAGAATCACAACCATCAGCAGATCATTCTAGACAAGGGCATGCTGGTGGCCATCGATGAGCTGATGCAGTCTCTGAATCATAATGGCGAGACTCTGCGCCAGAAACCTCCTGTGGGAGAAGCAGACCCTTACAGAGTGAAAATGAAGCTCTGCATCCTGCTTCACGCCTTCAGCACCCGCGTCGTGACCATCAACAGGGTGATGGGCTATCTGAGCTCCGCCTAA

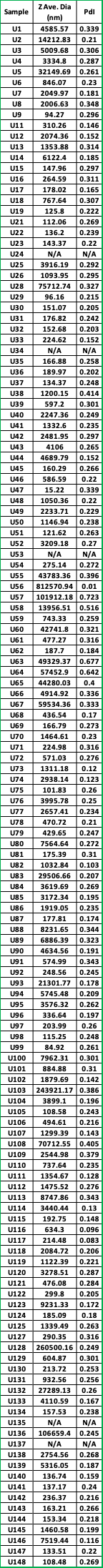

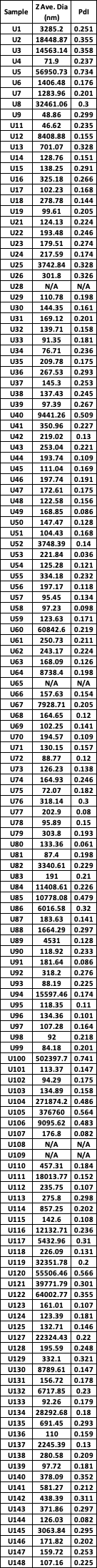

**Size by DLS:** average diameter and PDI of lipopolyplexes, loaded with Fluc mRNA in Tris-HCl buffer (pH 7.4)

**Size by DLS:** average diameter and PDI of lipopolyplexes, loaded with mCherry RNA in Tris-HCl buffer (pH 7.4)
